# Mutation Landscape of SARS COV2 in Africa

**DOI:** 10.1101/2020.12.20.423630

**Authors:** Angus A. Nassir, Clarisse Musanabaganwa, Ivan Mwikarago

## Abstract

COVID-19 disease has had a relatively less severe impact in Africa. To understand the role of SARS CoV2 mutations on COVID-19 disease in Africa, we analysed 282 complete nucleotide sequences from African isolates deposited in the NCBI Virus Database. Sequences were aligned against the prototype Wuhan sequence (GenBank accession: NC_045512.2) in BWA v. 0.7.17. SAM and BAM files were created, sorted and indexed in SAMtools v. 1.10 and marked for duplicates using Picard v. 2.23.4. Variants were called with mpileup in BCFtools v. 1.11. Phylograms were created using Mr. Bayes v 3.2.6. A total of 2,349 single nucleotide polymorphism (SNP) profiles across 294 sites were identified. Clades associated with severe disease in the United States, France, Italy, and Brazil had low frequencies in Africa (L84S=2.5%, L3606F=1.4%, L3606F/V378I/=0.35, G251V=2%). Sub Saharan Africa (SSA) accounted for only 3% of P323L and 4% of Q57H mutations in Africa. Comparatively low infections in SSA were attributed to the low frequency of the D614G clade in earlier samples (25% vs 67% global). Higher disease burden occurred in countries with higher D614G frequencies (Egypt=98%, Morocco=90%, Tunisia=52%, South Africa) with D614G as the first confirmed case. V367F, D364Y, V483A and G476S mutations associated with efficient ACE2 receptor binding and severe disease were not observed in Africa. 95% of all RdRp mutations were deaminations leading to CpG depletion and possible attenuation of virulence. More genomic and experimental studies are needed to increase our understanding of the temporal evolution of the virus in Africa, clarify our findings, and reveal hot spots that may undermine successful therapeutic and vaccine interventions.

## INTRODUCTION

SARS CoV2 virus is a positive-sense single stranded RNA(+ssRNA) coronavirus responsible for the covid-19 pandemic (Asghari *et al*., 2020). Since the initial isolation and genomic characterization of SARS CoV2 in January 2020, numerous mutation studies have tracked the evolution of the virus globally (Chaw *et al*., 2020; Korber *et al*., 2020; Koyama *et al*., 2020; Mishra *et al*., 2020; Tang *et al*., 2020; van Dorp *et al*., 2020). Mutation studies are important as they reveal important information about the temporal evolution of the virus, reveal suitable targets for drug, diagnostics, and vaccine design, and reveal hot spots that may undermine successful therapeutic and vaccine interventions (Kayla *et al*., 2018; Perales *et al*., 2011). Mutation studies also help track changes in the infectivity and virulence of mutants, reveal new strains with possible implications on immune escape and provide important clinico-epidemiological data (Abdullahi *et al*., 2020; Chen *et al*., 2020; Pachetti *et al*., 2020; Xi *et al*., 2020; Zou *et al*., 2020).

Studies have since shown that SARS CoV2 is a moderately mutating virus with a median mutation rate of 1.12 × 10−3 mutations per site-year (95% CI, CI: 9.86 × 10−4 to 1.85 × 10−4) (95% CI: 4.8 to 5.52) (Koyama *et al*., 2020). This moderate mutation rate is lower than that of other +ssRNA viruses and this is attributable to the presence of a 3’-5’ exonuclease that provides proof-reading ability (Duffy *et al*., 2018; Minskaia *et al*., 2006).

Mutation studies have also distinguished viral SARS CoV2 clades and mapped dominant strains. The six major SARS CoV2 clades include D614G or basal, L84S, L3606F, D448del and G392D. D614G was first seen in China and is now the dominant strain worldwide (Korber *et al*., 2020). The shift to the D614G variant occurs even in areas where the wild type strain is established and is due to the increased fitness of the mutant over the wild type (Korber *et al*., 2020).

Viral entry of SARS Cov2 is facilitated by cleavage of the SARS CoV2 S protein. The D614G mutation more efficiently facilitates its cleavage by the host serine protease elastase-2 and this explains the high infectivity of D614G (Hu *et al*., 2020). Increased infectivity of D614G is supported by *in vitro* experimental studies showing elevated RNA levels and higher viral titers in clinical samples with the D614G mutation and D614G mutant pseudoviruses respectively (Hu *et al*., 2020; Korber *et al*., 2020; Lorenzo-Redondo *et al*., 2020; Ozono *et al*., 2020; Wagner *et al*., 2020). However, there’s no conclusive evidence to show that the variant is associated with more severe disease or increased hospitalizations (Wagner *et al*., 2020). Even so, the D614G strain has co-evolved with other mutations such as (F106F), 14408 C->T (P323L), 241 C->T, 25563 G->T (Q57H), and 1059 C-> T(T85I) and more studies are required to clarify the impact of these mutations on virulence.

The D614G mutation is situated on the B cell epitope in a region that is highly immunodominant and this may possibly undermine vaccine effectiveness. However, experimental studies suggest that D614G mutants are sensitive to neutralization by polyclonal convalescent serum (Korber *et al*., 2020).

Subclades of D614G include D614G/Q57H/ and D614G/Q57H/T265I which were first identified in France, D614G/203_204delinsKR first identified in Germany and D614G/203_204delinsKR/T175M first identified in Iceland and Portugal (Koyama *et al*., 2020). The L84S clade was first observed in China and has one subclade namely L84S/P5828L that was first observed in the United States. The L3606F clade was also first observed in China and has the L3606F/V378I/ subclade first observed in Italy and the L3606F/G251V/ subclade observed in Brazil. Other subclades are D448del which was first observed in France and G392D which was first observed in Germany. In general, there’s high affinity between US and European samples with little similarity with East Asian samples and European clades dominate in samples in US (Koyama *et al*., 2020).

Mutations studies have also revealed the mechanisms of SARS CoV2 mutations. Dominant mutations are C->T transitions (Chaw *et al*., 2020; Koyama *et al*., 2020; Mishra *et al*., 2020). Depletion of CpG dinucleotides is also a common mutation mechanism in SARS CoV2. Increased CG dinucleotide levels are inversely correlated with viral fitness, defined by decreased virulence and replication. Thus, CpG depletion is defined as an immune escape strategy to evade host antiviral mechanisms. (Theys *et al*., 2018). Depletion of CpG dinucleotides in SARS CoV2 is possibly mediated by human zinc finger antiviral protein (hZAP) and apolipoprotein B mRNA editing enzyme (APOBEC1 and APOEBC3a). The hZAP attach to CpG dinucleotides in viral genomes to inhibit the replication of viruses and mediate the degradation of viral genome (Nchioua *et al*., 2020; Takata *et al*., 2017; Meagher *et al*., 2019; Trus *et al*., 2020). APOBEC1 and APOBEC3a deplete CpG dinucleotides in RNA viruses by mediating cytidine-to-uridine (C → T) changes (Xia, 2020; DiGeorgio *et al*, 2020).

In comparative terms, Africa has had lower covid-19 morbidity and mortality numbers. This is a striking observation especially when the vulnerabilities associated with weak or nonexistent health systems, poor sanitation, high HIV prevalence, and high poverty rates in Africa are taken into account (Anim & Ofor-Asenso, 2020; de Aranzabal *et al*., 2020; Patel *et al*., 2020). Despite this observation, and to the best of our knowledge, there have been no mutation studies focused on genomes sequenced from samples collected in Africa that seek to understand the evolution of this disease in Africa.

In this study, we analyzed SARS-CoV-2 genomes from 282 samples collected in Africa in order to characterize the genetic variants circulating in Africa and understand the virus’ temporal evolution. We evaluate if these mutations have clinically relevant outcomes, assessing implications on viral infectivity and disease severity in Africa and their potential effect on vaccine and therapeutic development and efficacy. We clarify the role of SARS CoV2 mutations in Covid-19 disease in Africa, relative to the rest of the world.

## MATERIALS AND METHODS

282 complete nucleotide sequences from African isolates were obtained from the NCBI Virus Database. The 282 sequences were aligned with the original Wuhan sequence (GenBank accession: NC_045512.2) (NCBI, 2020; Wang *et al*., 2020a) using the mem command in BWA v. 0.7.17 (Li, 2013). The SAM file was converted to BAM file using SAMtools v. 1.10 followed by sorting and indexing (Li *et al*., 2009). Duplicate marking and addition of read groups was done using Picard v. 2.23.4 (Broad Institute, 2019). Variants were called using the mpileup command in BCFtools v. 1.11 and visualized in IGV Viewer v. 2.8.9 (Robinson *et al*., 2011). Mutant proteins and their respective positions were abstracted from the called variants and the reference genome using a custom PHP script.

Multiple alignment was done in BLAST2 with the Bat coronavirus RaTG13 (GenBank accession: MN996532.2) as the outgroup (NCBI, 2020; Zheng *et al*., 2000). The BLAST2 dump file was converted into a Nexus file using the European Bioinformatics Institute (EBI) platform (Madeira *et al*., 2019). MEGA X: Molecular Evolutionary Genetics Analysis across computing platforms was used to select the general time reversible (GTR) evolutionary model (BIC score=340376.778) (Kumar, Stecher, Li, Knyaz, and Tamura 2018). Phylogenetic analysis of the aligned sequences involved maximum likelihood (ML) method in Mr. Bayes: Bayesian Inference of Phylogeny v 3.2.6 (ML lset nst=6, Lset rates=invgamma, parsimony model, default priors, ngen=10000, burning=82) (Huelsenbeck & Ronquist, 2001; Ronquist & Huelsenbeck, 2003).

The nexus translation tree was visualized using FigTree v 1.4.4 (Rambaut, 2010). The stability of point mutations was determined based on ddG values computed using I-Mutant 3.0, a support vector machine (SVM) tool that predicts protein stability upon single point mutations (Capriotti *et al*., 2005). ProtParam tool in the ExPasy server was used to compute thermal stability and other physicochemical properties of the mutated proteins (Gasteiger *et al*., 2005).

## RESULTS

A total of 282 nucleotide sequences from African isolates were analyzed. Majority of the sequences analyzed (80%) were from Egypt. 32.7% of sequences from the rest of Africa excluding Egypt were wild type with mutated sequences forming 67.3% of all non-Egyptian African sequences. All of the Egyptian sequences were mutated (Supplementary table S1).

70% (n=203) of mutations were transitions while 30% (n=86) were transversions. C-> T transitions formed 45% (n=130) of all mutations and 64% of transitions. G->T transversions were the second most dominant mutations and formed 20% (n=57) of all mutations and 66% of all transversions. T->G and C->G mutations were the least observed and when combined formed ∼1% of all mutations (n=2). 66% of all mutations (n=193) resulted in formation of T from other bases (table 1, figure 1).

**Table 1:** transition versus transversion mutations.

| TRANSITIONS |  | TRANSVERSIONS |  |
| --- | --- | --- | --- |
| C->T | 130 | C->A | 4 |
| T->C | 25 | A->C | 6 |
| A->G | 27 | G->T | 57 |
| G->A | 21 | T->G | 2 |
|  |  | A->T | 5 |
|  |  | G->C | 5 |
|  |  | T->A | 5 |
|  |  | C->G | 2 |
|  | 203 |  | 86 |

**Figure 1:**
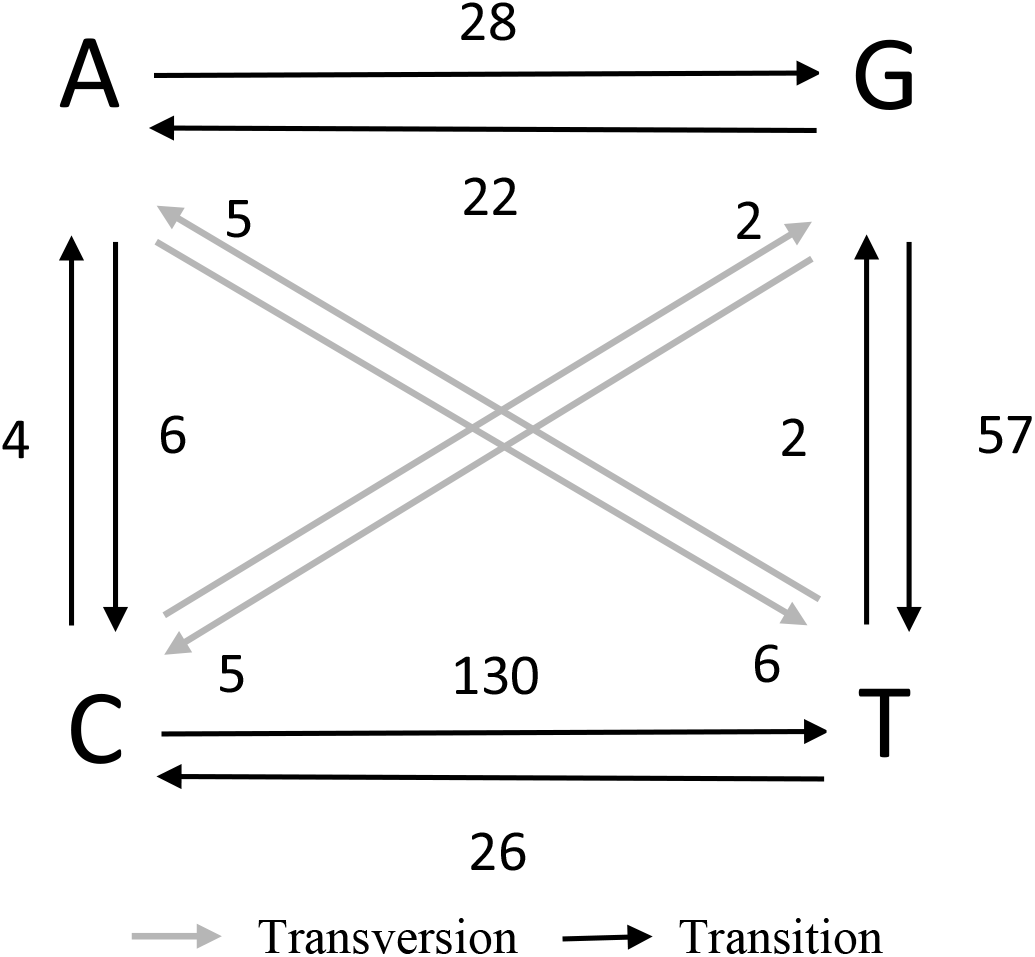
Transition / Transversion Diagrams showing SARS-CoV2 mutational patterns in Africa.

**Figure 2:**
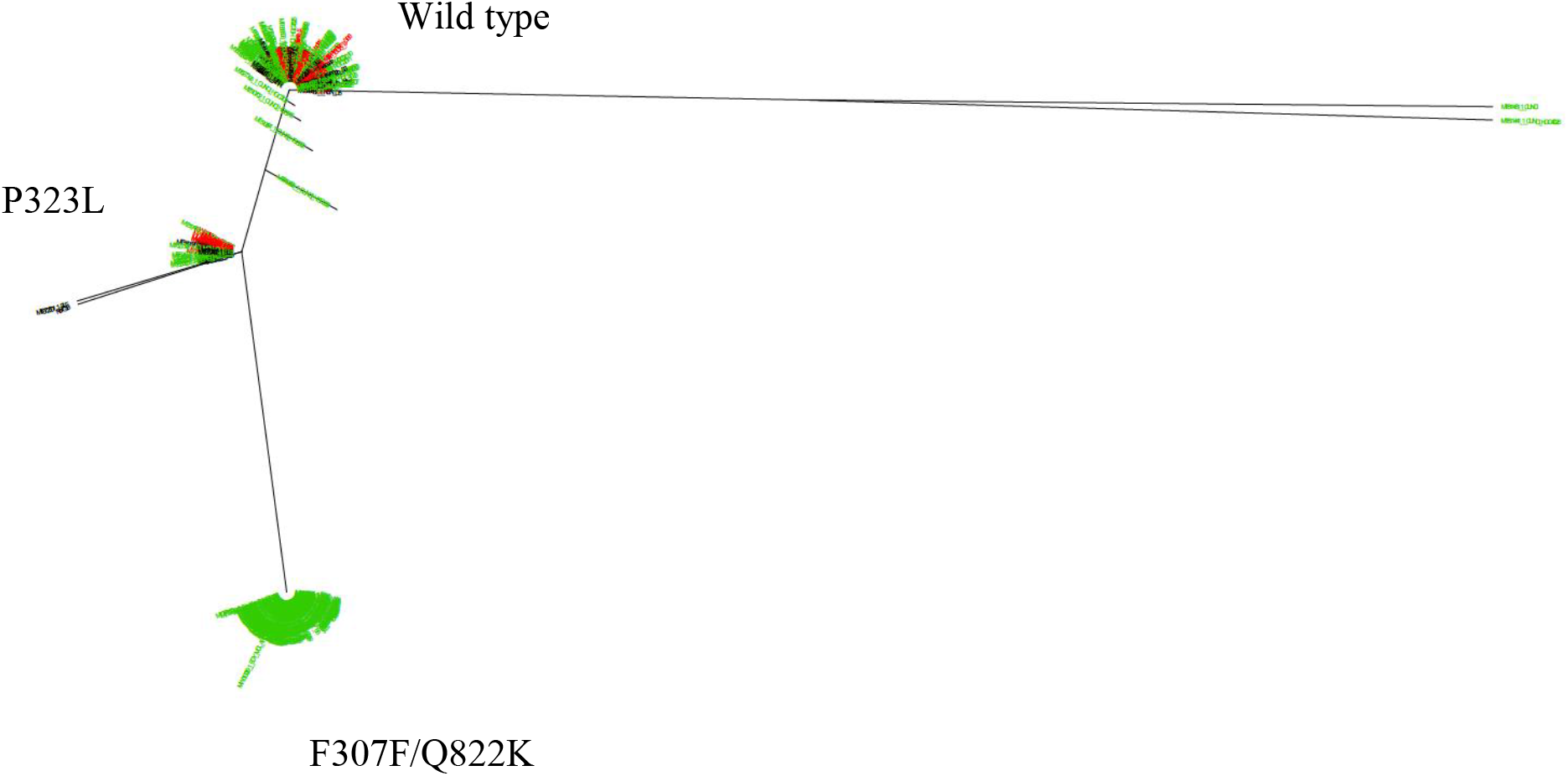
Phylogenetic tree constructed around the SARS CoV2 NSP12 region. Color coding was as follows: Egypt (green), Tunisia (purple), Morocco (orange), SSA (black). Samples from Italy (red) were also included. Samples are clustered into 3. The largest group was the F307F/Q822K which consisted of Egyptian samples only with F307F and Q822K always occurring together. P323L was well distributed in North Africa, occurred in a mutually exclusive fashion with F307F/822K, and consisted of only 3% of SSA samples. Majority of SSA samples were wild type. Italian samples were evenly distributed between the wild type and P323L groups.

For mutations with a frequency exceeding 1%, commonest mutations were missense mutations (54%) (table 2).

**Table 2:** SARS CoV2 mutations in Africa by type.

| Mutation type | Number | % total |
| --- | --- | --- |
| **Missense** | 107 | 54% |
| **Synonymous** | 72 | 37% |
| **Nonsense** | 1 | 1% |
| **Noncoding** | 17 | 9% |
| **Nonstop** | 0 | 0% |
|  | 197 | 100% |

A total of 4 indels were identified. All the indels were from Egyptian samples and affected the leader protein, RdRp, and the 3’ stem-loop II-like motif (s2m) respectively (Supplementary table S2).

The commonest SNPs were 3037 C->T (F106F), 23403 A->G (D614G), 14408 C->T (P323L), 241 C->T, 25563 G->T (Q57H), and 1059 C-> T(T85I). Common mutations such as 25563 G->T (Q57H), and 1059 C->T (T85I) were mostly present in samples from North Africa, specifically Morocco and Tunisia (Supplementary table S3).

98.7% of all Egyptian sequences had the 3037 C->T mutation (98.7%), 25403 A->G (98.2%), 241 C->T (95.2%), 25563 G->T (81.5%), 14362 C->T (63%), 29871 A->G (63%), 15907 G->A (62.6%), 17091 T->C (62.6%), and 26257 G->T (62.6%). The 14408 C->T (P323L) was not as widely observed in Egypt as it was in the rest of Africa (30% vs 70%). 14362C->T (L4788L), 29871A->G (noncoding), 15907G->A (Q822K), 17091T->C (G285G) and 26257G->T (V5F) accounted for more than half of mutations in Egyptian samples but were not observed in other African countries (Supplementary table S5).

### Non-Structural Protein Mutations

Majority of the mutations (∼63%) occurred in the NSP3 (16%), N (11.1%), S (11.1%), RdRp (9.9%), NSP4 (8%), and NSP2 (6.8%) regions respectively. Least mutated regions were s2m, ORF8, NSP16, Stem-loop 1, NSP10, and NSP7, each of which accounted for less than 1% of all mutations. Regions showing no mutations included NSP8, NSP9, NSP11, nsp15, ORF6, and ORF9, (Supplementary table S5). Majority of the mutations (>57%) in Egypt affected the nsp3 (17.8%), N nucleocapsid phosphoprotein (15.1%), S (13.8%), helicase (9.9%), and nsp2 (6%) regions respectively. Least mutated regions were ORF1ab/leader protein, ORF7b, E/envelope protein, ORF1ab/nsp10, ORF10/ Coronavirus 3’ UTR pseudoknot stem-loop 1 and ORF7a/ORF7a protein (Supplementary table S6).

### Structural Protein Mutations

Non-structural protein mutations exceeding 1% frequency occurred in the *E, N*, and *S* genes. More than half of all Egyptian samples (62.6%, n=141) had the 26257G->T (E, V5F) mutation and this was not observed in any other African country. The remaining 2 E mutations 26428 G->T(V62F) (1.06%) and 26299 C-> T(L19F) (0.35%) were present in Sierra Leone only (Supplementary table S8). The L19F mutation was characterized by an 8mer homopolymeric stretch (TTTTTTTT) and the V5F with a 4mer stretch (TTTT) (Supplementary table S7). Mutations on the *N* gene were geographically limited and all were observed in Tunisia except for 3. Egyptian samples had only 1 mutation on the nucleocapsid protein. The prevalence of 28881G->A(R203K), 28882G->A(R203R), and 28883G->C(G204R) was 8.5% for each mutation. 28878G->A(S202N) had a frequency of 2.48% (Supplementary table S8). M mutations were less than 1% in frequency.

The D614G mutation was the most dominant mutation on the spike glycoprotein. 249 (88.3%) of all sequences (n=282) had the D614G mutation. 47% (n=55) and 25% (n=6) of the non-Egyptian and non-North African samples respectively had the D614G mutation. Nearly all samples from Egypt (98%, n=227) had the D614G mutation. The D614G mutation was not observed in Kenyan and Zambian samples (figure 3, Supplementary table S9).

**Figure 3:**
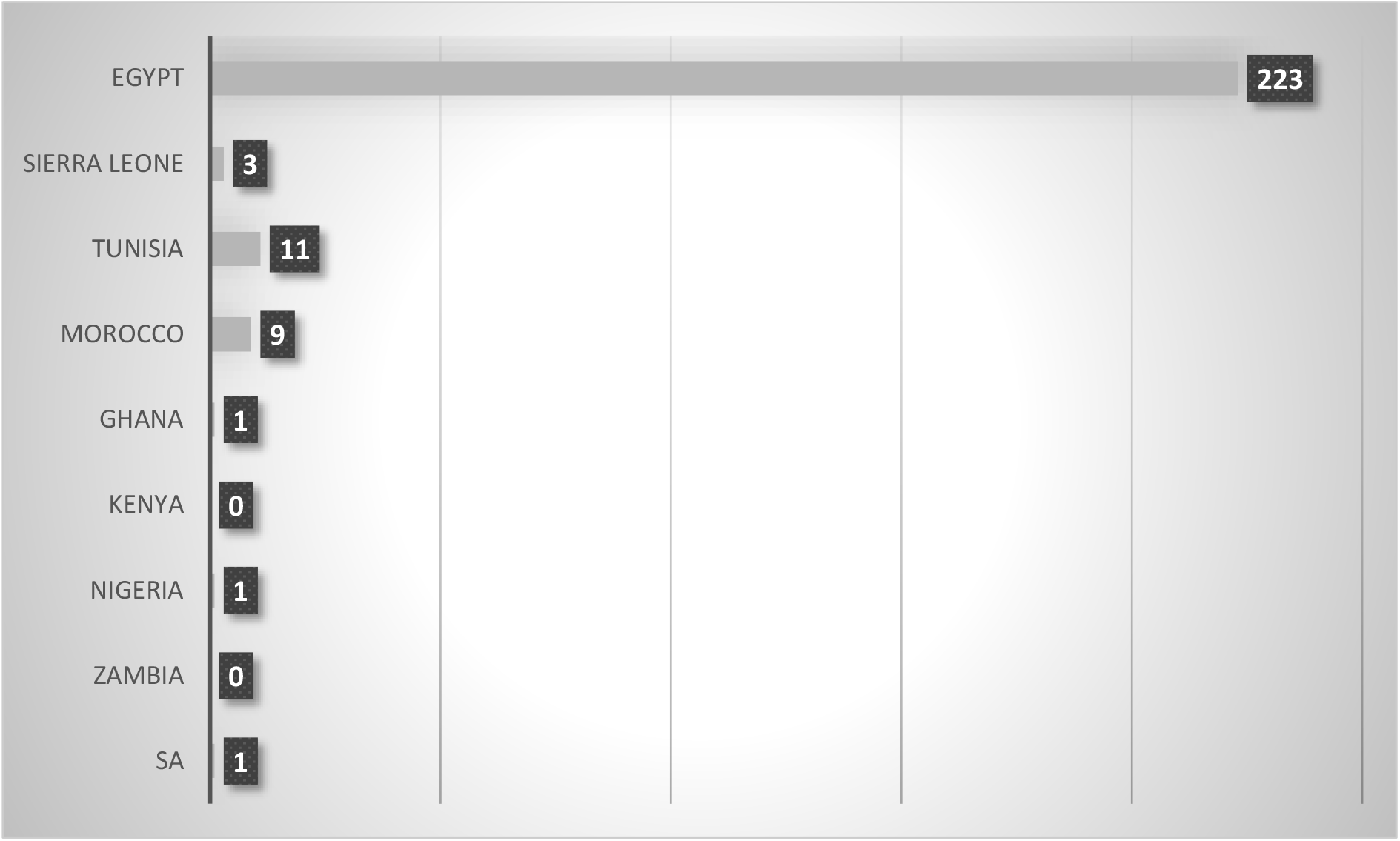
distribution of the SARS CoV2 D614G variant in early African samples.

In all samples, the D614G mutation co-occurred with 3037 C->T (F106F), 23403 A->G (D614G), 14408 C->T (P323L), and 241 C->T. The only exception to this rule was in Egypt where 14408 C->T (P323L) co-occurred with D614G in only 25% of all cases and 25563 G->T (Q57H) which co-occurred with D614G more than 71% all the time. 25563 G->K (Q57L) was observed in 4% of Egyptian samples. Co-occurrence of 14408 C->T (P323L) in samples from SSA was ∼3% (Supplementary table S27 and Supplementary table S34).

22468G->T(T302T) was the second most common spike glycoprotein mutation and was observed in 11% of all samples (Sierra Leone 27.2%, Tunisia 14.3%, Egypt 0.4%). 23127C->T(A522V) occurred on the spike GP receptor binding domain (RBD) was present in 9.5% of all Tunisian samples (Supplementary table S10). 69% were synonymous mutations, 31% missense mutations, 61.5% transition mutations and 38.5% transversion mutations.

### Structural Protein Mutations

Structural protein mutations are tabulated in Supplementary table S13-S25. Majority of the mutations occurred in the Nsp3 region. Structural protein mutations with an overall frequency greater than 1% included 1059C->T(T85I) on NSP2 (3.9%), 3037C->T(F106F) (88.7%) on NSP3, E310D on NSP4 (1.06%), and T148I on NSP8 (3.2%). 98.7% of Egyptian samples had T851 mutation compared to 47% for the rest of Africa. This was also the commonest synonymous NSP3 mutation worldwide (Koyama *et al*., 2020). A total of 16 different mutations were observed in all African samples. 14362 C->T (F307F) and 15907G->A(Q822K) had the highest frequencies, were observed in Egypt only and co-occurred together. The P323L mutation was less frequent in SSA (∼3%) and even though it was more widespread in North Africa, (∼30%), its frequency was lower in Egypt than the global frequency reported by Koyama *et al* (2020) (Supplementary table S21). 78.6% of all mutations were deamination mutations (C->T), 21.4% were G->T transversions. All resulted in the formation of T (Supplementary table S22). No mutations were observed for NSP10 and NSP11.

#### Accessory Protein Mutations

The commonest mutation of the ORF3a region was 25563G->T(Q57H). This mutation was prevalent in 71% of all African samples, with a frequency of 81.5% in Egypt, 60% in Morocco, 38.1% in Tunisia, and 9.1% in Sierra Leone. A unique mutation not observed elsewhere was Q57L, with a frequency of 4.4% in Egypt. Other mutations formed less than 1% of samples in Africa and these included 25411A->C(I7L), 26144G->T(G251V), and 25821C->T(A143A) (Supplementary table S26). Mutations on other accessory proteins had a frequency of less than 1% (figure 4, Supplementary table S27-30). No mutations were observed in the *ORF6* and *ORF10* genes.

**Figure 4:**
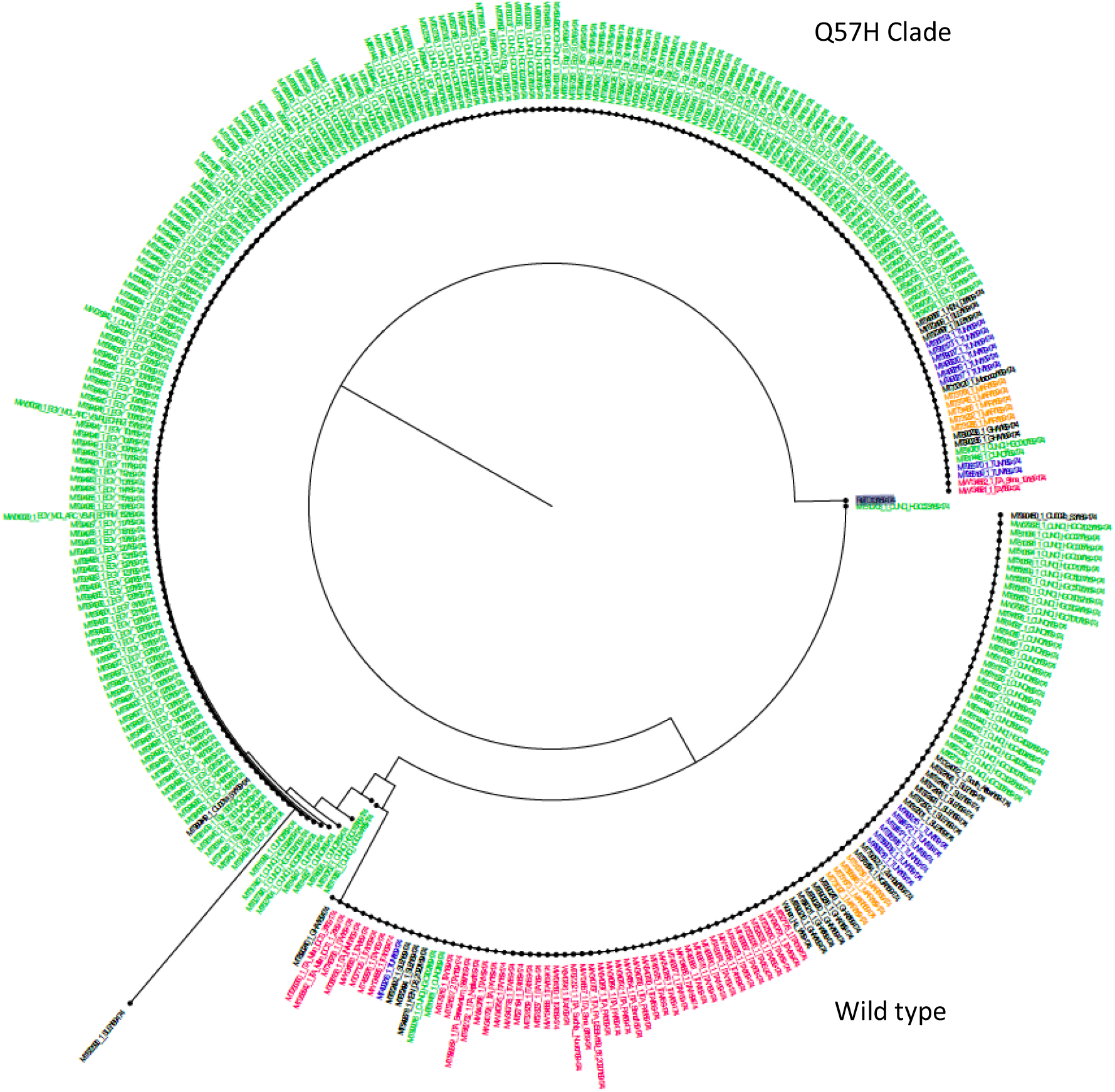
Phylogenetic tree constructed using the SARS CoV2 ORF3a region. Color coding was as follows: Egypt (green), Tunisia (purple), Morocco (orange), SSA (black). Samples from Italy (red) were also included. Samples are clustered into 2: one group had the Q57H mutation, another was wild type. Majority of Egyptian, Moroccan, and Tunisian samples belonged to the Q57H group. Majority of Italian and SSA samples were in the wild type group with only 4% of Italian samples in the Q57H group.

**Figure 5:**
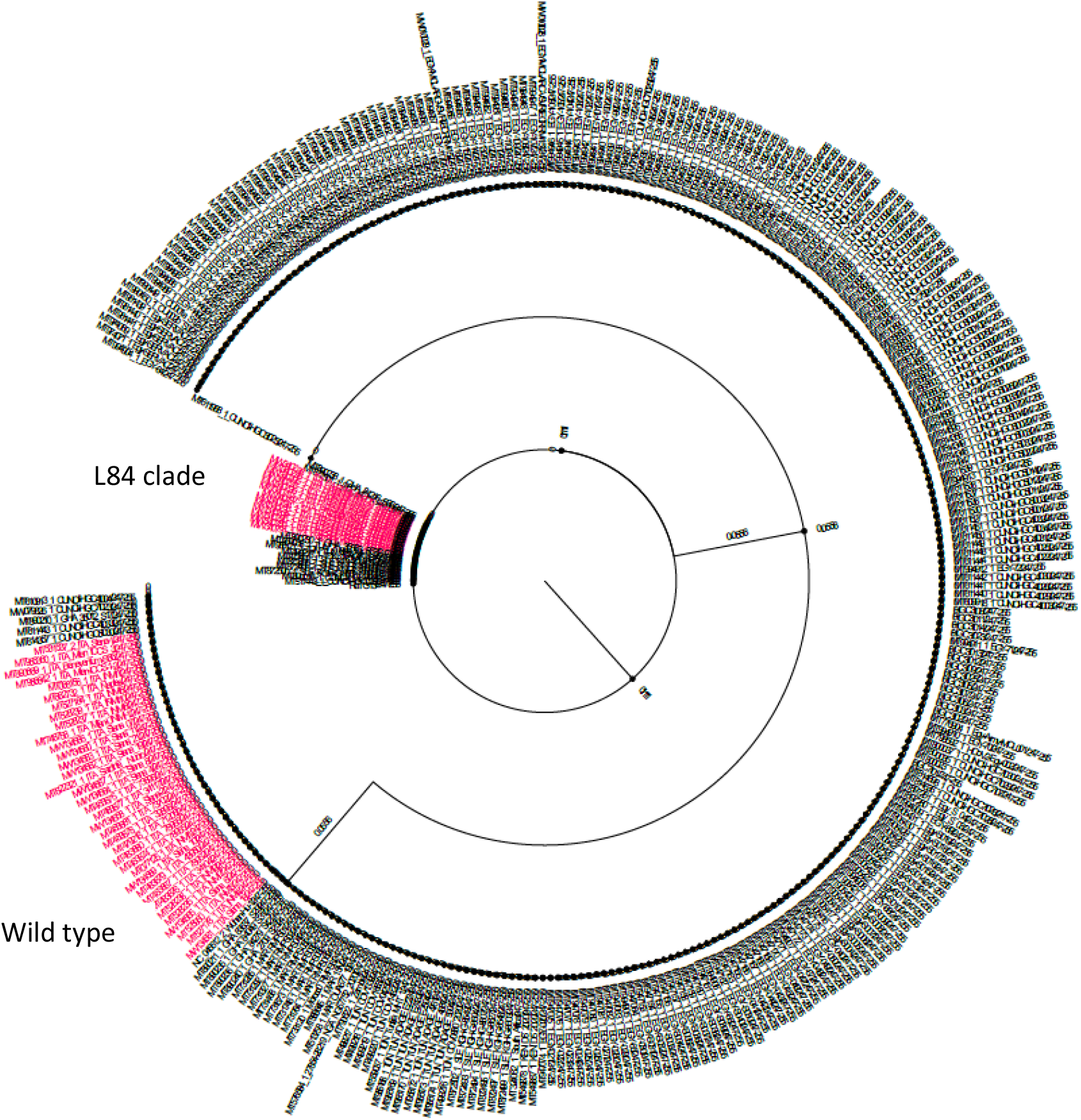
Phylogenetic tree constructed using the SARS CoV2 ORF8 region. Samples are clustered into 2: The L84S clade and the wild type. Italian samples (red) were included. Only 2.5% of African samples belonged to the L84S clade.

#### Regional Specificity

Mutations seen in Egypt and not observed elsewhere in Africa with a frequency of more than 1% included 15907G->A(Q822K) affecting the RdRp gene (62%), 26257G->T(V5F) affecting E (62.6%), 25563 G->K (Q57L) affecting ORF3a (4.4%), 10097G->A(G15S) affecting 3CLPro (4%), 12534C->T(T148I) affecting NSP8 (4%), and 28908G->T(G212V) affecting N (4%) (Supplementary table S31). A total of 3 missense and 2 synonymous mutations were observed in Moroccan samples only. The missense mutations occurred in 30% of the Moroccan samples and included 6404G->T(V1229F) affecting NSP3, 8208C->T(T1830I) affecting NSP3, and 29362C->T(P364H) affecting N (Supplementary table S31). 27.3% of samples from Sierra Leone had the 10818C->T(A255V) mutation affecting 3CLPro and not seen elsewhere in Africa (Supplementary table S32).

### Unreported Mutations

The following mutations were reported in other regions and not observed in Africa (table 3).

**Table 3:**
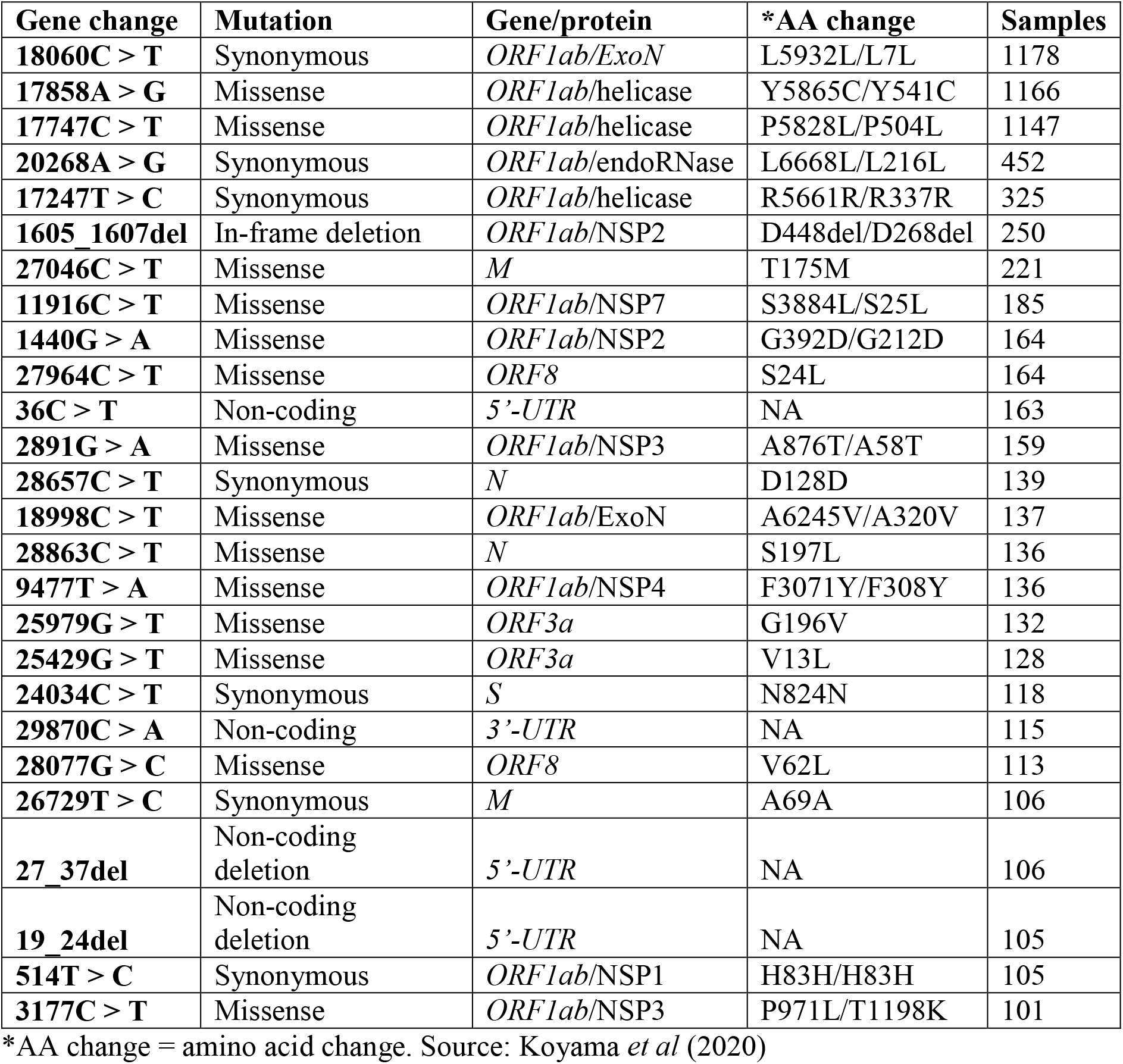
Mutations observed in other geographic regions but not observed in Africa.

| Gene change | Mutation | Gene/protein | *AA change | Samples |
| --- | --- | --- | --- | --- |
| **18060C > T** | Synonymous | *ORF1ab/ExoN* | L5932L/L7L | 1178 |
| **17858A > G** | Missense | *ORF1ab/helicase* | Y5865C/Y541C | 1166 |
| **17747C > T** | Missense | *ORF1ab/helicase* | P5828L/P504L | 1147 |
| **20268A > G** | Synonymous | *ORF1ab/endoRNase* | L6668L/L216L | 452 |
| **17247T > C** | Synonymous | *ORF1ab/helicase* | R5661R/R337R | 325 |
| **1605_1607del** | In-frame deletion | *ORF1ab/NSP2* | D448del/D268del | 250 |
| **27046C > T** | Missense | *M* | T175M | 221 |
| **11916C > T** | Missense | *ORF1ab/NSP7* | S3884L/S25L | 185 |
| **1440G > A** | Missense | *ORF1ab/NSP2* | G392D/G212D | 164 |
| **27964C > T** | Missense | *ORF8* | S24L | 164 |
| **36C > T** | Non-coding | *5'-UTR* | NA | 163 |
| **2891G > A** | Missense | *ORF1ab/NSP3* | A876T/A58T | 159 |
| **28657C > T** | Synonymous | *N* | D128D | 139 |
| **18998C > T** | Missense | *ORF1ab/ExoN* | A6245V/A320V | 137 |
| **28863C > T** | Missense | *N* | S197L | 136 |
| **9477T > A** | Missense | *ORF1ab/NSP4* | F3071Y/F308Y | 136 |
| **25979G > T** | Missense | *ORF3a* | G196V | 132 |
| **25429G > T** | Missense | *ORF3a* | V13L | 128 |
| **24034C > T** | Synonymous | *S* | N824N | 118 |
| **29870C > A** | Non-coding | *3'-UTR* | NA | 115 |
| **28077G > C** | Missense | *ORF8* | V62L | 113 |
| **26729T > C** | Synonymous | *M* | A69A | 106 |
| **27_37del** | Non-coding deletion | *5'-UTR* | NA | 106 |
| **19_24del** | Non-coding deletion | *5'-UTR* | NA | 105 |
| **514T > C** | Synonymous | *ORF1ab/NSP1* | H83H/H83H | 105 |
| **3177C > T** | Missense | *ORF1ab/NSP3* | P971L/T1198K | 101 |

\*AA change = amino acid change. Source: Koyama *et al* (2020)

## DISCUSSION

In the present study, we compared SARS-CoV-2 genomes from 282 samples collected in Africa against the Wuhan reference genome NC_045512.2 with the aim of understanding the evolution of the virus in Africa and the impacts of the mutations on morbidity and mortality in Africa.

We observed that 6% of all samples collected in Africa were wild type and 94% were mutants with 88% forming the D614G clade overall. 98% of all the Egyptian samples were the D614G variant and 98% of all D614G variants were from North Africa. If the Egyptian samples are excluded, wild type samples from the rest of Africa formed 34% of all sequences with the D614G clade comprising 25% of all SSA samples. This is in contrast to the earliest data demonstrating that the D614G variant formed 67% of all samples worldwide with higher running counts in Europe, Asia, Oceania, and North America and lower counts in parts of South America and Africa (Korber *et al*., 2020).

The wild type variant dominated the earliest samples collected from Kenya, Zambia, Ghana and Sierra Leone. Taken together with infection numbers, the data suggests that countries in which the earliest cases were caused by the wild type SARS CoV2 strain had relatively fewer infections. In converse, countries that had the highest number of infections in Africa such as South Africa, Morocco, Egypt, and Tunisia started off with a bigger proportion of the D614G variant. The predominance of the G614 variant in the early months of the pandemic in Africa may partly explain the relatively low numbers of those infected with covid-19 disease in many African countries. Predominance of the D614G variant during the early months of the pandemic may also explain the steep number of infections reported in South Africa and the North African countries of Egypt, Morocco, and Tunisia. This observation is based on findings that the D614G variant is associated with increased infectivity as clinical samples with the D614G mutation having higher viral titers (Hu *et al*., 2020; Korber *et al*., 2020; Lorenzo-Redondo *et al*., 2020; Ozono *et al*., 2020; Wagner *et al*., 2020). D614G is more infectious than the wild type sequence as it binds more efficiently to the human ACE2 receptor (Hu *et al*., 2020; Korber *et al*., 2020; Lorenzo-Redondo *et al*., 2020; Ozono *et al*., 2020; Wagner *et al*., 2020).

The WHO recently raised an alarm over the surge in infections in parts of Africa where the number of infections had remained low since the first case in Africa was reported in February. As pointed out by Korber *et al* (2020), the D614G variant has increased fitness and is under positive selection. Thus, the predominance may shift from the wild type to the D614G variant, allowing the latter to establish itself and predominate in areas where the wild type strain had been previously established with time (Korber *et al*., 2020; Koyama *et al*., 2020). Whereas there’s limited data of samples from Africa sequenced in the past few weeks, it is possible that the recent surge in cases in African countries that had hitherto low cases is due to the shift to the D614G variant. Genomic studies of current samples will help to clarify this.

Absence of other mutations on the spike glycoprotein appear to have influenced the course of the disease in Africa. Global studies have identified 4 other variants on the spike glycoprotein that appear to enhance virus pathogenicity. These variants are D364Y, V483A, G476S, and V367F all of which affect the S1 RBD domain. Ou *et al* (2020) observed that the V367F and D364Y variants confer more structural stability to the S protein and this enables the SARS CoV2 virus to bind more efficiently to the human ACE2 receptor (Ou *et al*., 2020). Experimental studies have demonstrated that V367F is associated with enhanced cell entry. Other RBD mutants that have been identified include N354D and W436R. N354D and D364Y, V367F, W436R had significantly lowered ΔG and significantly lowered equilibrium dissociation constant (KD) compared to the reference strain, suggesting that these mutants have significantly increased affinity to human ACE. In double mutants with N354D and D364Y, the latter provides increased affinity and this implies that the main contributor of the enhanced affinity is D364Y. Experimental validation assays prove that V367F significantly lowers the ED50 concentration of S and ACE2 receptor-ligand binding (Ou *et al*., 2020). These mutants are proposed to bind ACE2 more stably due to the enhancement of the base 208 rigidity (Ou *et al*., 2020). Our study did not identify any of these mutants in African samples and this may also explain the relatively lower morbidity and mortality seen in the African continent.

Another significant finding was on the occurrence of the P323L mutation on the RdRp protein. Globally, P323L is one of the commonest SARS CoV2 mutations with a frequency of over 90% in countries such as the United States (Koyama *et al*., 2020). In Africa however, the P323L mutation had an overall frequency of 34%, with only 3% of these mutations occurring in SSA. Whereas Egypt had a relatively high frequency of P323L (30.4%), only 25% of mutations co-occurred with D614G. Global data shows a near 100% correlation between D614G and P323L (Kannan *et al*., 2020).

P323L is thought to enhance viral infectivity together with D614G. According to Kannan *et al* (2020), the location of P323L on the NSP12-NSP8 interface may position the leucine side chain closer to F396, leading to enhanced hydrophobic interactions between NSP8 L122 residue and nsp12 T323 (Cγ2) and L270 residues. This is thought to enhance viral replication through improved processivity of NSP12 (Kannan *et al*., 2020). This observation seems to suggest that the low frequency of P323L in Africa may be a contributor to the relatively less severe impact of covid-19 disease in the continent compared to other regions even in areas such as Egypt where the D614G variant predominates.

We also observed the low frequency of the ORF8 L84S mutation in Africa. L84S was the first observed mutation and is one of the commonest mutation worldwide with a frequency exceeding 50% and associated with severe disease in Italy, (Koyama *et al*., 2020; Wang *et al*., 2020b). This mutation was present in 2.5% of African samples. The ORF8 protein aids in immune evasion through downregulation of major histocompatibility complex molecules class I (MCH-I) (Zhang *et al*., 2020). The L84S mutation is thought to decrease protein stability (DdG=0.99 kcal/mol) and protein rigidity, a factor that may disfavour SARS CoV2, leading to increased immune surveillance and reduced viral titres (Wang *et al*., 2020b). The effect of this mutation on disease severity and whether its low frequency in Africa contributes to the relatively low disease burden merits more experimental studies.

Other important findings related to the ORF3a protein. The first observation was that only 1 sample from SSA had the Q57H mutation on the ORF3a accessory protein. This mutation occurred in 81% of all Egyptian samples, in 60% of Moroccan samples and in 38% of Tunisian samples. The second observation was that the G251V ORF3a mutation which occurred first in Italy and Brazil and was associated with many infections formed less than 2% of all African samples (Koyama *et al*., 2020).

As reported by Koyama *et al*., Q57H is the commonest mutation worldwide (Koyama *et al*., 2020). Taken together with the earlier observations about the D614G variant, this observation suggests two things. First, it seems to point to Europe and or USA as the origin of the virus in much of North Africa since this the D614G/Q57H first occurred in France and has since then predominated in the USA (Koyama *et al*., 2020). Secondly, it may have implications on the disease burden in Africa. ORF3a interacts with both S and ORF8 proteins. According to Wu *et al* (2020), the Q57H mutation results in increased binding affinity between the Q57H Orf3a and S (ΔΔG = 4.2 kcal/mol). This dramatic increase in the binding affinity due to the Q57H mutation may have several consequences. First, it may cause failure of treatment by shifting the protein-binding interface and destroying drug-targeting sites (Wu *et al*., 2020). Secondly, it leads to formation of an early stop codon to orf3b after amino acid 13 (Δ3b), resulting in a truncated ORF3b protein with consequent loss of interferon antagonism (Lam *et al*., 2020). Other findings show that the Q57H variant does not seem to influence channel properties and does not result in any significant differences functionally compared to the wildtype ORF3a. This may be attributed to the mutation being located on the N-terminal which determines the subcellular localization of the virus without influencing channel properties (Kern *et al*., 2020). Further research on the clinical importance of Q57H is warranted.

In the present study, we also observed 3 missense mutations in the E protein: V5F, V62F, and L19F that may be of clinic-epidemiological importance. E is a 75-residue integral viroporin involved in viral replication, pathogenesis and assembly, activation of host inflammasome, and virion release (Lim & Liu, 2001; Nieto-Torres *et al*., 2014; Ruch & Machamer, 2012; Weiss & Navas-Martin, 2005). E is highly conserved in coronaviruses with very few observed mutations (Qingfu *et al*., 2003). Deletion of E is associated with attenuation in some coronaviruses. Reduced virulence due to E mutations has been reported (De Diego *et al*., 2007; Nieto-Torres *et al*., 2014; Pervushin *et al*., 2009). E is hence a suitable target for drug and vaccine development and channel activity may be optimally inhibited by targeting small-molecule drugs to host cell Golgi and the endoplasmic reticulum–Golgi intermediate compartment (ERGIC) (Mandala *et al*., 2020).

Structurally, E is made up of an N-terminal domain (NTD), an ion-conducting transmembrane domain (TMD), and a cytoplasmic domain (CTD) (Wu *et al*., 2003; Mandala *et al*., 2020). The V5F mutation affects the NTD and was present in more than half of all Egyptian samples (62.6%, n=141). This mutation has not been reported elsewhere to the best of our knowledge. The V19F and V62F mutations affect the TMD and CTD respectively. These 2 mutations are characterized by 8mer (TTTTTTTT) and 4mer (TTTT) homopolymeric stretches. Mutation analysis using the I-Mutant Suite indicate DdG values of 0.77 Kcal/mol, −1.21 Kcal/mol, and - 1.04 Kcal/mol for V62F, L19F and V5F. This suggests that the mutations result in highly unstable and temperature-sensitive E proteins. This observation is consistent with the work of Fischer *et al* (1998) who investigated the effect of E mutations on reduced thermostability and morphology aberrance (Fischer *et al*., 1998).

Since the E protein is essential for induction of interferon synthesis and apoptosis, RNA replication, and production and release of membrane vesicles or virus-like particles (VLPs) in coronaviruses (An *et al*., 1999; Corse & Machamer, 2000; Maeda *et al*., 2001; De Diego *et al*., 2007; Nieto-Torres *et al*., 2014; Pervushin *et al*., 2009; Mandala *et al*., 2020; Wu *et al*., 2003), the observed mutations may impact the pathogenicity of SARS CoV2 in Africa. The mutations may also hinder effective detection of the virus using currently available RT-PCR kits based on amplification of the E protein. As noted, ∼63% of samples from Egypt had the V5F mutation. It is not clear if this mutation has an impact on disease severity and currently available RT-PCR and other diagnostic tests for SARS CoV2. More studies are needed to clarify the effect of these mutations on viral pathogenicity and viral detection using currently available RT-PCR kits.

Observations about the mutations on the E protein need also to be discussed in conjunction with ORF3a since ORF3a is also a viroporin-coding gene in coronaviruses (An *et al*., 1999; Jiang *et al*., 2005; Verdia-Baguena *et al*., 2012). In SARS CoV2, ORF3a forms homotetrameric potassium sensitive ion channels (viroporin) that mediates the activation of NLRP3 inflammasome (Siu *et al*., 2019; Farag *et al*., 2020; Wozniak *et al*., 2010). Viroporin subunits undergo oligomerization, forming hydrophilic pores that allow ions to be shuttled across the membranes of host cells and facilitate the cellular entry of viruses and release of viruses from infected cells and viral replication and assembly (Farag *et al*., 2020). Deletion of genes coding for viroporins leads to a significant reduction in viral progeny formation and reduces the pathogenicity of viruses (Farag *et al*., 2020). As noted previously, viroporins induce inflammaosme activity. Inflammasomes can regulate the activity of caspase-1. Caspase-1 mediates interleukin-1 β (IL-1β) and interleukin 18 (IL-18) maturation. In turn, IL-1β and IL-18 (Farag *et al*., 2020).

E and ORF3a proteins are thought to contribute to NLRP3 inflammasome activity and are essential for maximal replication and virulence of SARS CoV (Farag *et al*., 2020). SARS CoV viruses lacking E and ORF3a are not viable. Even though it contributes to viral pathogenesis, ORF3a in SARS CoV is not essential for replication (Siu *et al*., 2020).

Siu *et al* (2019) showed that ORF3a-associated activation of NLRP3 inflammasome activity is mediated through TNF receptor-associated factor 3 (TRAF3)–mediated ubiquitination of apoptosis-associated speck-like protein containing a caspase recruitment domain (ASC) (Siu *et al*., 2019). Findings by Siu *et al* (2020) also demonstrate that ORF3a up-regulates expression of fibrinogen subunits FGA, FGB and FGG in host lung epithelial cells in SARS CoV. ORF3a is also involved in the induction of apoptosis in cell culture and in the downregulation of type 1 interferon receptor through induction of serine phosphorylation within the IFN alpha-receptor subunit 1 (IFNAR1) degradation motif and increasing the ubiquitination of IFNAR1 (Siu *et al*., 2019). Based on the foregoing, the effect of the E and ORF3a mutations merit further investigations.

Majority (>70%) of the mutations were transition mutations, with C->T transitions making up 44% of these transitions. This finding is similar to observations in other studies that show dominance of C->T mutations in SARS CoV2 genome (Koyama *et al*., 2020; Mishra *et al*., 2020; Wang *et al*., 2020b; Badua *et al*., 2020). G->T transversions were the second most common mutations in our study followed by A->G and T->C transitions. Together, C->T and G->T transitions made up 64% of all mutations, underlining the role of deamination in SARS CoV2 evolution.

CpG depletion is an important mechanism deployed by RNA viruses to evade host antiviral proteins. This is because unmethylated CpG dinucleotides stimulate TLR 9 innate immune responses. Since CpG-rich codons have a lower transcription rate, CpG depletion also serves to enhance the virus transcription rate. Depletion of CpG dinucleotides may occur in response to selection pressure from host immune system, from spontaneous deamination of methylated cytosines in CpG dinucleotdes, and deamination of unmethylate d cytosines. Depletion of CpG is also a strategy that ensures the epigenetic silencing of the virus, leading to establishment of latent viral infection (Bird, 1980; Chinnery *et al*., 2012; Medvedeva *et al*., 2010; Wiebauer *et al*., 1993). Depletion may be driven by host ZAP or APOBEC proteins. In the present study, we observed that 95% of all the mutations in the NSP12 (RdRp) protein are deamination mutations. The role of RdRp deamination mutations in attenuating viral virulence in SARS CoV2 needs to be investigated further.

Findings reported in the present study may have clinico-epidemiological implications. We have noted the absence or presence in low frequencies of mutations associated with increased infectivity, virus fitness, and disease severity. Experimental studies will help to clarify the clinical impact of these mutations. On vaccines, the RBD has important epitopic antigens. Some of the identified mutations may alter the binding affinity of vaccines raised against the prototype strain hence leading to a reduction in vaccine efficacy (Ou *et al*., 2020). Further studies also need to investigate if the CpG depletion that is widespread on the SARS CoV2 RdRp is a strategy to attenuate viral virulence (Ficarelli *et al*., 2020; Trus *et al*., 2020) and immune escape and determine if it has a role in the relatively low morbidity and mortality numbers in Africa.

Mutation analysis is a critical factor when selecting suitable targets for drug design. Currently, there are a number of drugs in use, undergoing clinical trials, or proposed as suitable drug targets against SARS CoV2 genomic or sub-genomic RNA regions. Lopinavir and ritonavir were proposed for repurposing of SARS CoV2 3-chymotrypsin-like (3CLpro) and papain-like (PLpro) proteases (Nutho *et al*., 2020; Xiaopan *et al*., 2020). Remdesivir and Favipravir target the RdRp (Goldman *et al*., 2020; Pandey *et al*., 2020) while Ribavirin interferes with mRNA capping and viral replication (Khalili *et al*., 2020; Pandey *et al*., 2020; Tong *et al*., 2020). Osetalmivir which targets the 3CLPro was found to be ineffective in the treatment of SARS CoV2 (Tan *et al*., 2020). Interferons induce production of Mx proteins that are thought to inhibit viral replication (Spiegel *et al*., 2004). Teicoplanin is thought to prevent viral entry through inhibition of cathepsin L (Zhang *et al*., 2020). Baricitinib which was discovered using artificial intelligence (AI) methods, prevents viral entry by binding to AP2-associated protein kinase 1 (AAK1) to block clathrin-dependent endocytosis and modulation of inflammatory cytokines through selective inhibition of Janus Kinase (JAK) (Caputo *et al*., 2020). Other compounds that have been mentioned as possible drugs for covid-19 disease include azithromycin (Echeverría-Esnal *et al*., 2020) and arbidol (Gao *et al*., 2020). Except for remdesivir and corticosteroids, studies on many of these other drugs are still ongoing or have produced conflicting results (Echeverría-Esnal *et al*., 2020; Pandey *et al*., 2020).

Mutations may have an impact on the binding of SARS CoV2 antiviral drugs and this has implications on drug design and drug resistance. This has been noted for antiviral drugs targeting the RdRp regions and the effectiveness of compounds such as remdesivir may be hampered by the mutations (Pandey *et al*., 2020). On drug design, regions such as PLpro, RdRp, and S that exhibited high mutation rates may be less suitable drug and diagnostics targets, and regions showing limited or no mutations such as ORF8, NSP8, NSP9, NSP11, nsp15, ORF6, 2’-O-ribose methyltransferase, and ORF9 may be more attractive targets. Assessment of viral mutations provided protein stability data and can be used to model protein folding as well as assess binding affinity around the mutation. This helps to assess possible drug resistance phenotypes, select optimal targets for lead candidate development, correlate mutations with disease severity, and map putative target sites. Further studies need to consider these mutations in this respect.

### Limitations of the Study

Out of the 18,820 global SARS CoV2 sequences deposited in NCBI, African sequences accounted for 280 or just 1.5% of all sequences and Egypt accounts for 80% of these sequences collected in Africa at the time of the analysis. Sub-Saharan Africa accounted for 0.29% of all sequences. Only 9 African countries had some SARS CoV2 sequences; not a single sequence was seen for 45 other countries. Evidently, very little effort is being made to sequence samples collected in Africa and understand the mutation patterns in this continent. The small sample size may not be sufficient to make sweeping generalizations. The genetic picture captured in this study is a temporal screenshot that explains the genetic variation present months ago. The mutation landscape is a constantly changing mosaic that is temporal in nature and which requires constant genomic analysis for continuous tracking.

