## Supplementary Information for "Mutation Landscape of SARS COV2 in Africa"

### SUPPLEMENTARY FILES

***Table S1: Number of sequences analyzed by country and the proportion of wild-type versus mutated sequences***

| COUNTRY | TOTAL SEQS | WILDTYPE | | MUTATED | NUMBER OF MUTATIONS |
|---|---|---|---|---|---|
| ***Kenya** | 2 | 1 | 50.00% | 50.00% | 6 |
| **Zambia** | 1 | 1 | 100.00% | 0.00% | 0 |
| **Nigeria** | 1 | 0 | 0.00% | 100.00% | 9 |
| **SA** | 1 | 0 | 0.00% | 100.00% | 8 |
| **Ghana** | 8 | 5 | 62.50% | 37.50% | 21 |
| **Morocco** | 10 | 1 | 10.00% | 90.00% | 74 |
| **Tunisia** | 21 | 6 | 28.57% | 71.43% | 176 |
| **Sierra Leone** | 11 | 4 | 36.36% | 63.64% | 64 |
| **Egypt** | 227 | 0 | 0.00% | 100.00% | 1994 |
| **TOTAL** | **282** | **18** | **6.38%** | **93.62%** | **2352** |

TOTAL SEQS=total sequences; *incomplete sequences.

***Table S2: indel mutations observed in African SARS CoV2 isolates***

| POS | REF | ALT | GENE | PROTEIN AFFECTED |
|---|---|---|---|---|
| **685** | AAAGTCATTT | A | ORF1ab | Leader protein |
| **14474** | GATACCACTTCAGAGAGCTAGGTGTTGTA | GA | RdRp | RdRp |
| **14486** | GAGAGCTAGGTGTTGTA | GA | RdRp | RdRp |
| **29735** | AGGCCACGCGGAGTACGATCGAGTGTACAGTG | A | | 3' stem-loop II-like motif (s2m) |

POS=genomic position; REF=wild type nucleotide; ALT=mutated nucleotide;

***Table S3: Table Sof the commonest SARS-CoV2 SNPs from African isolates.***

| POS | REF | ALT | SA | ZAM | NGR | KEN | GH | MOR | TUN | SL | TOTAL | % TS | % MS |
|---|---|---|---|---|---|---|---|---|---|---|---|---|---|
| **3037** | C | T | 1 | 0 | 1 | | 1 | 9 | 11 | 3 | 26 | 47.27% | 70.27% |
| **23403** | A | G | 1 | 0 | 1 | | 1 | 9 | 11 | 3 | 26 | 47.27% | 70.27% |
| **14408** | C | T | 1 | 0 | 1 | | 1 | 9 | 11 | 3 | 26 | 47.27% | 70.27% |
| **241** | C | T | 1 | 0 | 1 | | 1 | 9 | 10 | 3 | 25 | 45.45% | 67.57% |
| **25563** | G | T | | | | | | 6 | 8 | 1 | 15 | 27.27% | 40.54% |
| **1059** | C | T | | | | | | 5 | 4 | 1 | 10 | 18.18% | 27.03% |
| **8782** | C | T | | | | | 1 | | 3 | 3 | 7 | 12.73% | 18.92% |
| **28144** | T | C | | | | | 1 | | 3 | 3 | 7 | 12.73% | 18.92% |
| **28878** | G | A | | | | | 1 | | 3 | 3 | 7 | 12.73% | 18.92% |
| **22468** | G | T | | | | | | | 3 | 3 | 6 | 10.91% | 16.22% |
| **29742** | G | A | | | | | 1 | | 2 | 3 | 6 | 10.91% | 16.22% |
| **28881** | G | A | | | 1 | | 1 | | 2 | 1 | 5 | 9.09% | 13.51% |
| **28882** | G | A | | | 1 | | 1 | | 2 | 1 | 5 | 9.09% | 13.51% |
| **28883** | G | C | | | 1 | | 1 | | 2 | 1 | 5 | 9.09% | 13.51% |
| **18877** | C | T | | | | | | 1 | 3 | | 4 | 7.27% | 10.81% |
| **41** | T | C | | | | | | | 4 | | 4 | 7.27% | 10.81% |
| **11083** | G | T | | | | 1 | | | 2 | 1 | 4 | 7.27% | 10.81% |
| **15324** | C | T | | | | | | 3 | 1 | | 4 | 7.27% | 10.81% |

POS=genomic position; REF=wild type nucleotide; ALT=mutated nucleotide; SA=South Africa; ZAM=Zambia; NGR=Nigeria; KEN=Kenya; GH=Ghana; MOR=Morocco; TUN=Tunisia; SL=Sierra Leone; TS=Sequences without Egypt; MS=Total sequences including Egypt.

*Table S4: Commonest Mutations in Egypt*

| POS | REF | ALT | EGYPT | % TOTAL |
|---|---|---|---|---|
| **3037** | C | T | 224 | 98.68% |
| **23403** | A | G | 223 | 98.24% |
| **241** | C | T | 216 | 95.15% |
| **25563** | G | T | 185 | 81.50% |
| **14362** | C | T | 143 | 63.00% |
| **29871** | A | G | 143 | 63.00% |
| **15907** | G | A | 142 | 62.56% |
| **17091** | T | C | 142 | 62.56% |
| **26257** | G | T | 142 | 62.56% |
| **14408** | C | T | 69 | 30.40% |
| **18877** | C | T | 25 | 11.01% |
| **28881** | G | A | 19 | 8.37% |
| **28882** | G | A | 19 | 8.37% |
| **28883** | G | C | 19 | 8.37% |
| **29744** | G | A | 12 | 5.29% |
| **28849** | C | T | 11 | 4.85% |

POS=genomic position; REF=wild type nucleotide; ALT=mutated nucleotide; SA=South Africa; ZAM=Zambia; NGR=Nigeria; KEN=Kenya; GH=Ghana; MOR=Morocco; TUN=Tunisia; SL=Sierra Leone; TS=Sequences without Egypt; MS=Total sequences including Egypt.

*Table S5: Gene/protein regions affected by mutations in Africa excluding Egypt*

| *Gene* | Protein | Number | % |
|---|---|---|---|
| *ORF1ab* | NSP3 (papain like protease) | 26 | 16.05% |
| *N* | Nucleocapsid phosphoprotein | 18 | 11.11% |
| *S* | Surface Glycoprotein | 18 | 11.11% |
| *ORF1ab* | NSP12 (RdRp) | 16 | 9.88% |
| *ORF1ab* | NSP4 | 13 | 8.02% |
| *ORF1ab* | NSP2 | 11 | 6.79% |
| *5’-UTR* | Noncoding | 8 | 4.94% |
| *3’-UTR* | Noncoding | 6 | 3.70% |
| *M* | Membrane Glycoprotein | 5 | 3.09% |
| *ORF3a* | ORF3a protein | 5 | 3.09% |
| *ORF1ab* | NSP5 (3C-like proteinase) | 5 | 3.09% |
| *ORF1ab* | NSP15 (endoRNAse) | 4 | 2.47% |
| *ORF1ab* | NSP14 (3'-to-5' exonuclease) | 4 | 2.47% |
| *ORF1ab* | NSP1 (leader protein) | 4 | 2.47% |
| *E* | Envelope protein | 3 | 1.85% |
| *ORF1ab* | NSP13 (Helicase) | 3 | 1.85% |
| *ORF1ab* | NSP6 | 3 | 1.85% |
| *ORF7b* | ORF7b protein | 2 | 1.23% |
| *ORF7a* | ORF7a protein | 2 | 1.23% |

| | | | |
|---|---|---|---|
| *s2m* | | 1 | 0.62% |
| *ORF8* | ORF8 protein | 1 | 0.62% |
| *ORF1ab* | NSP16 (2'-O-ribose methyltransferase) | 1 | 0.62% |
| *ORF1ab* | Stem-loop 1 | 1 | 0.62% |
| *ORF1ab* | NSP10 | 1 | 0.62% |
| *ORF1ab* | NSP7 | 1 | 0.62% |
| | | **162** | **100.00%** |

***Table S6: Mutations by gene/protein region in Egypt***

| GENE /PROTEIN | NUMBER | % |
|---|---|---|
| ORF1ab/nsp3 | 27 | 17.76% |
| N/ nucleocapsid phosphoprotein | 23 | 15.13% |
| S/ surface glycoprotein | 21 | 13.82% |
| ORF1ab/helicase | 15 | 9.87% |
| ORF1ab/nsp2 | 9 | 5.92% |
| ORF1ab/RdRp | 7 | 4.61% |
| ORF1ab/nsp4 | 7 | 4.61% |
| ORF1ab/ 3'-to-5' exonuclease | 7 | 4.61% |
| 3’UTR | 5 | 3.29% |
| ORF3a/ ORF3a protein | 4 | 2.63% |
| ORF8/ORF8 protein | 4 | 2.63% |
| ORF1ab/3C-like proteinase | 3 | 1.97% |
| ORF1ab/nsp8 | 3 | 1.97% |
| ORF1ab/2'-O-ribose methyltransferase | 3 | 1.97% |
| M/membrane glycoprotein | 2 | 1.32% |
| ORF1ab/nsp6 | 2 | 1.32% |
| ORF1ab/nsp7 | 2 | 1.32% |
| ORF6/ORF6 protein | 2 | 1.32% |
| ORF1ab/leader protein | 1 | 0.66% |
| ORF7b | 1 | 0.66% |
| E/envelope protein | 1 | 0.66% |
| ORF1ab/nsp10 | 1 | 0.66% |
| ORF10/ Coronavirus 3' UTR pseudoknot stem-loop 1 | 1 | 0.66% |
| ORF7a/ORF7a protein | 1 | 0.66% |
| | **152** | **100.00%** |

***Table S7: Mutations in the envelope protein of SARS CoV 2 samples from Africa. Three missense mutations were observed in the envelope protein. The prevalence of 26428 G->T (V62F) was 27.3% while that of 26299 C->T(L19F) was in 9.1% of all Sierra Leonean samples.***

| **POS** | **REF** | **ALT** | **MUTATION** | **TYPE** | **COUNTRY** | **FREQ** |
|---|---|---|---|---|---|---|
| 26428 | G | T | V62F | Missense | Sierra Leone | 27.3% |
| 26299 | C | T | L19F | Missense | Sierra Leone | 9.1% |
| 26257 | G | T | V5F | Missense | Egypt | 62.6% |
| **Synonymous** | | | 0 | **Transition** | 1 | 33.3% |
| **Missense** | | | 3 | **Transversion** | 2 | 66.7% |
| **Nonstop** | | | 0 | | | |

POS=genomic position; REF=wild type nucleotide; ALT=mutated nucleotide; FREQ=Frequency.

***Table S8: Mutations in the N nucleocapsid protein of SARS CoV 2 samples from Africa. 28878G->A(S202N) had a frequency of 12.5% in Ghana, 14.3% in Tunisia, 27.3% in Sierra Leone, and 0.4% in Egypt***

| **POS** | **REF** | **ALT** | **MUTATION** | **TYPE** | **COUNTRY** | **FREQ** | **EGYPT** |
|---|---|---|---|---|---|---|---|
| 28878 | G | A | S202N | Missense | GH, TUN, SL | 12.7% | 0.35% |
| 28881 | G | A | R203K | Missense | NG, TUN, SL, GH | 9.1% | 8.4% |
| 28882 | G | A | R203R | Missense | NG, TUN, SL, GH | 9.1% | 8.4% |
| 28883 | G | C | G204R | Missense | NG, TUN, SL, GH | 9.1% | 8.4% |
| 28311 | G | C | P13L | Missense | Sierra Leone | 0.35% | 0.0% |
| 28346 | G | T | G25C | Missense | Tunisia | 0.35% | 0.0% |
| 28347 | G | T | G25V | Missense | Tunisia | 0.35% | 0.0% |
| 28531 | C | T | Y86Y | Synonymous | Tunisia | 0.35% | 0.0% |
| 28688 | T | C | L139L | Synonymous | Tunisia | 0.35% | 0.0% |
| 28738 | T | A | A155A | Synonymous | Morocco | 0.35% | 0.0% |
| 28830 | C | A | S186Y | Missense | Tunisia | 0.35% | 0.0% |
| 28835 | T | C | S188P | Missense | Tunisia | 0.35% | 0.0% |
| 28854 | C | T | S194L | Missense | Tunisia | 0.35% | 0.0% |
| 28985 | G | T | G238C | Missense | Tunisia | 0.35% | 0.0% |
| 29315 | G | C | D348H | Missense | Morocco | 0.35% | 0.0% |
| 29449 | G | T | V393V | Missense | Tunisia | 0.35% | 0.0% |
| 29474 | G | T | D401Y | Missense | Tunisia | 0.35% | 0.0% |
| **Synonymous** | | 3 | 17.6% | | | | |
| **Missense** | | 14 | 82.4% | | | | |
| **Transition** | | 11 | 64.7% | | | | |
| **Transversion** | | 6 | 35.3% | | | | |

POS=genomic position; REF=wild type nucleotide; ALT=mutated nucleotide; SA=South Africa; ZAM=Zambia; NGR=Nigeria; KEN=Kenya; GH=Ghana; MOR=Morocco; TUN=Tunisia; SL=Sierra Leone; TS=Sequences without Egypt; MS=Total sequences including Egypt.

*Table S9:distribution of the D614G mutation by country*

| | D614G Strains | Total Sequences | Percentage |
|---|---|---|---|
| **Kenya** | 1 | 2 | 50.00% |
| **Zambia** | 0 | 1 | 0.00% |
| **Nigeria** | 1 | 1 | 100.00% |
| **SA** | 0 | 1 | 0.00% |
| **Ghana** | 1 | 8 | 12.50% |
| **Morocco** | 9 | 10 | 90.00% |
| **Tunisia** | 11 | 21 | 52.38% |
| **Sierra Leone** | 3 | 11 | 27.27% |
| **Egypt** | 223 | 227 | 98.24% |
| **Total** | **249** | **282** | **88.30%** |
| **Non-Egyptian** | 26 | 55 | 47.27% |

*Table S10: Mutations in the spike glycoprotein of SARS CoV 2 samples from Africa*

| POS | REF | ALT | MUTATION | TYPE | LOCATION | COUNTRY | FREQ |
|---|---|---|---|---|---|---|---|
| 23403 | A | G | D614G | Missense | S1 | All except Kenya | 47.30% |
| 22468 | G | T | T302T | Synonymous | NTD | Tunisia, SL | 11.00% |
| 23127 | C | T | A522V | Missense | RBD | Tunisia | 3.60% |
| 22397 | T | A | Y279N | Missense | NTD | Tunisia | 1.80% |
| 22424 | C | T | A288V | Missense | NTD | Tunisia | 1.80% |
| 22435 | T | C | C291R | Missense | NTD | Tunisia | 1.80% |
| 22503 | A | G | Q314Q | Synonymous | RBD | Tunisia | 1.80% |
| 23923 | C | T | Q787H | Missense | S2 | SL | 1.80% |
| 24370 | C | T | S937L | Missense | S2, HR 1 | Ghana | 1.80% |
| 24926 | G | T | V1122V | Synonymous | S2 | Ghana | 1.80% |
| 24992 | G | C | E1144E | Synonymous | S2 | SL | 1.80% |
| 25217 | G | T | G1219C | Missense | S2 | Ghana | 1.80% |
| 21575 | C | T | L5F | Missense | SP | Ghana, Egypt | 1.80% |

POS=genomic position; REF=wild type nucleotide; ALT=mutated nucleotide; S1=Spike Protein 1; NTD=N-terminal domain; RBD=receptor-binding domain; S2=Spike protein 2; HR1= heptapeptide repeat sequence 1; SP=signal peptide; FREQ=percentage frequency in Africa excluding Egypt.

***Table S11: Mutations in the membrane glycoprotein of SARS CoV 2 samples from Africa. 60% of membrane glycoprotein mutations were synonymous mutations. 26665T->C(I48T) was observed in a single Kenyan sequence and 26775G->T(A85S) in 4.8% of Tunisian samples (Table S11).***

| POS | REF | ALT | MUTATION | TYPE | COUNTRY | FREQ |
|---|---|---|---|---|---|---|
| 26665 | T | C | I48T | Missense | Kenya | 1.80% |
| 26720 | G | C | V66V | Synonymous | Tunisia | 1.80% |
| 26775 | G | T | A85S | Missense | Tunisia | 1.80% |
| 26790 | T | C | L90L | Synonymous | Tunisia | 1.80% |
| 26801 | C | G | L93L | Synonymous | Sierra Leone | 1.80% |

POS=genomic position; REF=wild type nucleotide; ALT=mutated nucleotide; FREQ=percentage frequency in Africa excluding Egypt.

*Table S12: mutations observed in the leader protein of SARS CoV2 samples in Africa*

| POS | REF | ALT | MUTATION | TYPE | COUNTRY | FREQ |
|---|---|---|---|---|---|---|
| 315 | G | T | S17I | Missense | Sierra Leone | 1.82% |
| 391 | A | G | A42A | Synonymous | Egypt | 0.35% |
| 745 | C | T | N160N | Synonymous | Ghana | 1.82% |

POS=genomic position; REF=wild type nucleotide; ALT=mutated nucleotide; S1=Spike Protein 1; NTD=N-terminal domain; RBD=receptor-binding domain; S2=Spike protein 2; HR1= heptapeptide repeat sequence 1; SP=signal peptide; FREQ=percentage frequency in Africa/Egypt.

*Table S13: NSP2 mutations observed in SARS CoV2 samples in Africa*

| POS | REF | ALT | MUTATION | TYPE | COUNTRY | FREQ |
|---|---|---|---|---|---|---|
| 1059 | C | T | T85I | Missense | Mor, Tun, SL, Egypt | 3.90% |
| 2113 | C | T | I436I | Synonymous | Tunisia, Egypt | 1.06% |
| 1404 | C | T | P200L | Missense | Tunisia, Egypt | 0.71% |
| 1191 | C | T | P129L | Missense | Kenya | 0.35% |
| 1397 | G | A | V198I | Missense | Tunisia | 0.35% |
| 1666 | T | C | F288F | Synonymous | Tunisia | 0.35% |
| 2306 | C | T | L501F | Missense | Ghana | 0.35% |
| 2416 | C | T | Y537Y | Synonymous | Tunisia | 0.35% |
| 2480 | A | G | I559V | Missense | Kenya | 0.35% |
| 2558 | C | T | P585S | Missense | Kenya | 0.35% |

POS=genomic position; REF=wild type nucleotide; ALT=mutated nucleotide; S1=Spike Protein 1; NTD=N-terminal domain; RBD=receptor-binding domain; S2=Spike protein 2; HR1= heptapeptide repeat sequence 1; SP=signal peptide; FREQ=percentage frequency in Africa excluding Egypt.

***Table S14: mutations observed in NSP3 region of SARS CoV2 samples in Africa.***

| POS | REF | ALT | MUTATION | TYPE | COUNTRY | FREQ |
|---|---|---|---|---|---|---|
| 3037 | C | T | F106F | Synonymous | All except Keya and Zambia | 88.7% |
| 5284 | C | T | N855N | Synonymous | Egypt | 3.2% |
| 3373 | C | A | D218E | Missense | Egypt | 2.5% |
| 4002 | C | T | T428I | Missense | Egypt | 2.1% |
| 5020 | C | T | D767D | Synonymous | Egypt | 2.1% |
| 3011 | T | C | L98L | Synonymous | Egypt | 1.8% |
| 4354 | G | T | E545G | Missense | Egypt | 1.8% |
| 2841 | C | T | A41V | Missense | Egypt | 1.1% |
| 3340 | G | T | V207V | Synonymous | Tunisia | 0.4% |
| 3896 | G | T | V393F | Missense | Tunisia | 0.4% |
| 4288 | G | T | E523G | Missense | Tunisia | 0.4% |
| 5359 | T | C | A880A | Synonymous | Tunisia | 0.4% |
| 5401 | A | G | A894A | Synonymous | Ghana | 0.4% |
| 5557 | T | C | G946V | Missense | Tunisia | 0.4% |
| 5672 | C | T | P985S | Missense | Ghana | 0.4% |
| 5986 | C | T | F1089F | Synonymous | Ghana | 0.4% |
| 6040 | C | T | F1107F | Synonymous | Sierra Leone | 0.4% |
| 6312 | C | A | T1198L | Missense | Sierra Leone | 0.4% |
| 6337 | G | T | S1206S | Synonymous | Tunisia | 0.4% |
| 7279 | C | T | F1520F | Synonymous | Tunisia | 0.4% |
| 7420 | C | T | I1567I | Synonymous | Tunisia | 0.4% |
| 7765 | C | T | S1682S | Synonymous | Tunisia | 0.4% |
| 8097 | C | T | T1793I | Missense | Tunisia | 0.4% |
| 8371 | G | T | Q1884H | Missense | Tunisia | 0.4% |
| 8389 | C | T | N1890N | Synonymous | Tunisia | 0.4% |

POS=genomic position; REF=wild type nucleotide; ALT=mutated nucleotide; S1=Spike Protein 1; NTD=N-terminal domain; RBD=receptor-binding domain; S2=Spike protein 2; HR1= heptapeptide repeat sequence 1; SP=signal peptide; FREQ=percentage frequency in Africa / Egypt.

***Table S15: mutations observed in the NSP4 protein of SARS CoV2 samples in Africa. Missense mutations with the highest frequencies included S34F, S76N and G309C which are found in Tunisia with a frequency of 4.8%. Overall frequencies however were below 1%. ound in Egypt only had a frequency of 1.06% (Table S14).***

| POS | REF | ALT | MUTATION | TYPE | COUNTRY | FREQ |
|---|---|---|---|---|---|---|
| 8655 | C | T | S34F | Missense | Tunisia | 0.35% |
| 8781 | G | A | S76N | Missense | Tunisia | 0.35% |
| 8950 | C | T | I132I | Synonymous | Sierra Leone | 0.35% |
| 9479 | G | T | G309C | Missense | Tunisia | 0.35% |
| 9514 | A | G | L320L | Synonymous | Tunisia | 0.35% |
| 9214 | C | T | Y220Y | Synonymous | Tunisia | 0.71% |
| 9483 | A | C | E310D | Missense | Egypt | 1.06% |
| 9520 | C | T | F322F | Synonymous | Egypt | 1.06% |
| 10046 | G | A | V498I | Missense | Tunisia, Egypt | 0.71% |

POS=genomic position; REF=wild type nucleotide; ALT=mutated nucleotide; S1=Spike Protein 1; NTD=N-terminal domain; RBD=receptor-binx jding domain; S2=Spike protein 2; HR1= heptapeptide repeat sequence 1; SP=signal peptide; FREQ=percentage frequency.

***Table S16: mutations observed in the NSP5 region of SARS CoV2 samples in Africa. NSP5 (3C-like proteinase, 3CLpro). Top missense NSP5 mutations were A255V and R279C which were present in 27.3% of Sierra Leonean samples and in 9.5% of Tunisian samples respectively but made 1% or less of all African samples***

| POS | REF | ALT | MUTATION | TYPE | | COUNTRY | FREQ |
|---|---|---|---|---|---|---|---|
| **10456** | C | T | F134F | Synonymous | Transition | Tunisia | 0.35% |
| **10582** | C | T | D176D | Synonymous | Transition | Tunisia | 0.35% |
| **10818** | C | T | A255V | Missense | Transition | Sierra Leone | 1.06% |
| **10889** | C | T | R279C | Missense | Transition | Tunisia | 0.71% |
| **10097** | G | A | G15S | Synonymous | Transition | Egypt | 3.2% |

***Table S17: mutations observed in the NSP6 region of SARS CoV2 samples in Africa. NSP6***

***A single missense NSP6 mutation, L37F, was observed in Kenya, Tunisia and Sierra Leone. Total frequency of this mutation was 1.4% (7.3% excluding Egyptian samples)***

| POS | REF | ALT | MUTATION | TYPE | COUNTRY | FREQ |
|---|---|---|---|---|---|---|
| 11083 | G | T | L37F | Missense | Kenya, Tunisia, SL | 1.4% |
| 11575 | C | T | F201F | Synonymous | Tunisia | 0.35% |
| 11782 | A | G | K270K | Synonymous | SL | 0.35% |

***Table S18: mutations observed in the NSP7 region of SARS CoV2 samples in Africa. NSP7 Two missense mutations on NSP7 were observed with an overall frequency of 0.35% (E50G, 1.4% in Egypt) and Q31H (4.8% in Tunisia) (Table S17).***

| POS | REF | ALT | MUTATION | TYPE | COUNTRY | FREQ |
|---|---|---|---|---|---|---|
| 11991 | A | G | E50G | Missense | Egypt | 1.06% |
| 11935 | A | C | Q31H | Missense | Tunisia | 0.35% |

POS=genomic position; REF=wild type nucleotide; ALT=mutated nucleotide; FREQ=percentage frequency in Africa.

***Table S19: mutations observed in the NSP8 region of SARS CoV2 samples in Africa***

| POS | REF | ALT | MUTATION | TYPE | COUNTRY | FREQ |
|---|---|---|---|---|---|---|
| 12534 | C | T | T148I | Missense | Egypt | 3.2% |

POS=genomic position; REF=wild type nucleotide; ALT=mutated nucleotide; FREQ=percentage frequency in Africa.

***Table S20: mutations observed in the NSP9 region of SARS CoV2 samples in Africa***

| POS | REF | ALT | MUTATION | TYPE | COUNTRY | FREQ |
|---|---|---|---|---|---|---|
| 12970 | C | T | N95N | Synonymous | Morocco | 1.06% |

POS=genomic position; REF=wild type nucleotide; ALT=mutated nucleotide; FREQ=percentage frequency in Africa.

***Table S21: mutations observed in the NSP12 region of SARS CoV2 samples in Africa***

| POS | REF | ALT | MUTATION | TYPE | COUNTRY | FREQ |
|---|---|---|---|---|---|---|
| 14362 | C | T | F307F | Synonymous | Egypt | 63% |
| 15907 | G | A | Q822K | Missense | Egypt | 62.60% |
| 14408 | C | T | P323L | Missense | SA, NG, GH, MOR, TUN, SL, EGY | 47.30% |
| 15324 | C | T | N628N | Synonymous | Morocco, Tunisia | 7.30% |
| 13536 | C | T | Y32Y | Synonymous | Egypt | 2.50% |
| 13515 | C | T | G25G | Synonymous | Ghana | 1.80% |
| 13517 | C | T | T26I | Missense | Ghana | 1.80% |
| 13620 | C | T | D60D | Synonymous | South Africa | 1.80% |
| 13730 | C | T | A97V | Missense | Sierra Leone | 1.80% |
| 13858 | G | T | D140Y | Missense | Tunisia | 1.80% |
| 14184 | C | T | T248T | Synonymous | Kenya | 1.80% |
| 14805 | C | T | Y455Y | Synonymous | Kenya | 1.80% |
| 15372 | G | T | T644T | Synonymous | Tunisia | 1.80% |
| 15380 | G | T | S647I | Missense | Morocco | 1.80% |
| 16178 | C | T | S913L | Missense | Tunisia | 1.80% |
| 14805 | C | T | Y455Y | Synonymous | Kenya | 1.80% |

POS=genomic position; REF=wild type nucleotide; ALT=mutated nucleotide; FREQ=percentage frequency in Africa.

***Table S22: RdRp mutation types and frequencies***

| Mutation Type | Number | Percentage | Mutation Type | Percentage |
|---|---|---|---|---|
| Synonymous | 9 | 56.2% | Transition | 81.2% |
| Missense | 7 | 43.8% | Transversion | 18.8% |
| | | | C->T transitions | 75% |
| | | | G->T transitions | 18.8% |
| | | | G->A transversions | 6.2% |
| **TOTAL** | **16** | 100.0% | Mutations resulting to T | 93.8% |

***Table S23: mutations on the NSP13 region of SARS CoV2 in Africa. NSP13 (Helicase). 4 synonymous mutations were observed, the most dominant of which was 17091T->C(G285G) with a prevalence of 50.4%. 16512A->G(L92L) had a 27.3% prevalence in Sierra Leone but just 1% overall frequency (Table S22).***

| POS | REF | ALT | MUTATION | TYPE | COUNTRY | FREQ |
|---|---|---|---|---|---|---|
| 17550 | C | T | L438L | Synonymous | Egypt | 1.1% |
| 17766 | C | T | V510V | Synonymous | Egypt | 1.1% |
| 16512 | A | G | L92L | Synonymous | Sierra Leone | 1.06% |
| 17091 | T | C | G285G | Synonymous | Egypt | 50.35% |

POS=genomic position; REF=wild type nucleotide; ALT=mutated nucleotide; FREQ=percentage frequency in Africa.

***Table S24: mutations on the NSP14 region of SARS CoV2 in Africa. All the mutations are observed in samples from the North African countries of Morocco, Tunisia, and Egypt. NSP14 (3'-5' exoribonuclease). Mutations on the NSP14 region included 18508C->T(L157F) with a prevalence of 10% in Morocco, 18928C->T(P297S) with a prevalence of 4.8% in Tunisia, and 19525G->T(D496Y) with a prevalence of 4.8% in Tunisia. Overall, all these formed less than 1% of all mutations in Africa and were missense mutations. A synonymous mutation, 18877C->T(L280L), had an overall prevalence of 10.3% and was present in North Africa (Table S23).***

| POS | REF | ALT | MUTATION | TYPE | COUNTRY | FREQ |
|---|---|---|---|---|---|---|
| 18508 | C | T | L157F | Missense | Morocco | 0.35% |
| 18928 | C | T | P297S | Missense | Tunisia | 0.35% |
| 19525 | G | T | D496Y | Missense | Tunisia | 0.35% |
| 18877 | C | T | L280L | Synonymous | Morocco, Tunisia, Egypt | 10.3% |

***Table S25: mutations on the NSP15 region of SARS CoV2 in Africa. NSP15 (endoRNAase / A1 and nsp15B-NendoU). Three missense mutations on the NSP15 region were observed. 19735G->T(D39V), 20032C->T(R138C), and 20374A->G(I252V) had an overall prevalence of 0.35% in Africa (Table S24).***

| POS | REF | ALT | MUTATION | TYPE | COUNTRY | FREQ |
|---|---|---|---|---|---|---|
| 19735 | G | T | D39V | Missense | Sierra Leone | 0.35% |
| 20032 | C | T | R138C | Missense | Tunisia | 0.35% |
| 20374 | A | G | I252V | Missense | Nigeria | 0.35% |

POS=genomic position; REF=wild type nucleotide; ALT=mutated nucleotide; FREQ=percentage frequency in Africa.

***Table S26: one missense mutation was observed in the Ribose 2'-O-methyltransferase region of SARS CoV2 samples in Africa. NSP16 (Ribose 2'-O-methyltransferase/ 2'-o-MT). One missense mutation, 20759C->T(A34V) was observed in Tunisia with an overall frequency of 0.35% in Africa. the frequency of the mutation in Tunisia was 4.8% (Table S25).***

| POS | REF | ALT | MUTATION | TYPE | COUNTRY | FREQ |
|---|---|---|---|---|---|---|
| 20759 | C | T | A34V | Missense | Tunisia | 4.8% |

POS=genomic position; REF=wild type nucleotide; ALT=mutated nucleotide; FREQ=percentage frequency in Africa.

***Table S27: mutations observed in the ORF3a region of SARS CoV2 samples in Africa***

| POS | REF | ALT | MUTATION | TYPE | COUNTRY | FREQ |
|---|---|---|---|---|---|---|
| 26144 | G | T | G251V | Missense | Egypt | 1.1% |
| 25563 | G | T | Q57H | Missense | Egypt | 70.9% |
| 25563 | G | K | Q57L | Missense | Egypt | 4.4% |
| 25821 | C | T | A143A | Synonymous | Sierra Leone | 0.71% |
| 25411 | A | C | I7L | Missense | Tunisia | 0.35% |

POS=genomic position; REF=wild type nucleotide; ALT=mutated nucleotide; FREQ=percentage frequency in Africa.

***Table S28: mutations observed in the ORF7a region of SARS CoV2 samples in Africa. One nonsense mutation 27754G->T(Z121END) was observed in Tunisia. The overall frequency of this mutation in Africa was 0.35% and its frequency in Tunisia was 4.8%. 27520A->T(N43Y) was observed in Sierra Leone with a frequency of 9.1% in Sierra Leone and an overall frequency of 0.35% in Africa (Table S27).***

| POS | REF | ALT | MUTATION | TYPE | COUNTRY | FREQ |
|---|---|---|---|---|---|---|
| 27520 | A | T | N43Y | Missense | Sierra Leone | 4.8% |
| 27754 | G | T | Z121END | Nonsense | Tunisia | 9.1% |

POS=genomic position; REF=wild type nucleotide; ALT=mutated nucleotide; FREQ=percentage frequency in Africa.

***Table S29: mutations observed in the OR7b region of SARS CoV2 samples in Africa. A single missense mutation was observed in ORF7b with an overall frequency of 0.35% (Table S30).***

| POS | REF | ALT | MUTATION | TYPE | COUNTRY | FREQ |
|---|---|---|---|---|---|---|
| 27769 | C | T | S5L | Missense | Tunisia | 0.35% |
| 27804 | C | T | L17L | Synonymous | Tunisia | 0.35% |

POS=genomic position; REF=wild type nucleotide; ALT=mutated nucleotide; FREQ=percentage frequency in Africa.

***Table S30: mutations observed in the ORF8 region of SARS CoV2 samples in Africa. There was a single mutation 28144T->C(L84S) with a frequency of 0.35% in Egypt (Table S29).***

| POS | REF | ALT | MUTATION | TYPE | COUNTRY | FREQ |
|---|---|---|---|---|---|---|
| 28144 | T | C | L84S | Missense | Egypt | 0.35% |

POS=genomic position; REF=wild type nucleotide; ALT=mutated nucleotide; FREQ=percentage frequency in Africa.

***Table S31: Mutations observed only in Egypt and not in other African countries***

| POSITION | REF | ALT | AA CHANGE | REGION | FREQUENCY |
|---|---|---|---|---|---|

| | | | | | | |
|---|---|---|---|---|---|---|
| 14362 | C | T | F307F | RdRp | 143 | 63.00% |
| 29871 | A | G | Noncoding | 3'-UTR | 143 | 63.00% |
| 15907 | G | A | Q822K | RdRp | 142 | 62.56% |
| 17091 | T | C | G285G | Helicase | 142 | 62.56% |
| 26257 | G | T | V5F | Envelope | 142 | 62.56% |
| 29744 | G | A | Noncoding | 3'-UTR | 12 | 5.29% |
| 28849 | C | T | N192N | N | 11 | 4.85% |
| 5284 | C | T | N855N | NSP3 | 9 | 3.96% |
| 10097 | G | A | G15S | 3CLPro | 9 | 3.96% |
| 12534 | C | T | T148I | NSP8 | 9 | 3.96% |
| 28908 | G | T | G212V | N | 9 | 3.96% |
| 23731 | C | T | T723T | S | 8 | 3.52% |
| 3373 | C | A | D218E | NSP3 | 7 | 3.08% |
| 4002 | C | T | T428I | NSP3 | 7 | 3.08% |
| 13536 | C | T | Y32Y | RdRp | 7 | 3.08% |
| 5020 | C | T | D767D | NSP3 | 6 | 2.64% |
| 3011 | T | C | L98L | NSP3 | 5 | 2.20% |
| 4354 | G | T | E545G | NSP3 | 5 | 2.20% |
| 28846 | C | T | R191R | N | 5 | 2.20% |
| 2841 | C | T | A41V | NSP3 | 3 | 1.32% |
| 9483 | A | C | Y220Y | NSP4 | 3 | 1.32% |
| 9520 | C | T | E310D | NSP4 | 3 | 1.32% |
| 11991 | A | G | E50G | NSP7 | 3 | 1.32% |
| 17550 | C | T | L438L | Helicase | 3 | 1.32% |
| 17766 | C | T | V510V | Helicase | 3 | 1.32% |
| 21597 | C | T | S12F | S | 3 | 1.32% |

POSITION=genomic position; REF=wild type nucleotide; ALT=mutated nucleotide; AA Change=amino acid change; FREQUENCY=percentage frequency in Egypt.

***Table S32: Mutations observed only in Morocco and not in other parts of Africa***

| **POSITION** | **REF** | **ALT** | **AA CHANGE** | **REGION** | **NUMBER** | **FREQUENCY %** |
|---|---|---|---|---|---|---|
| 6404 | G | T | V1229F | *NSP3* | 3 | 30% |
| 8208 | C | T | T1830I | NSP3 | 3 | 30% |
| 12970 | C | T | N95N | NSP9 | 3 | 30% |
| 22363 | G | T | V801V | S | 3 | 30% |
| 29362 | C | T | P364H | N | 3 | 30% |

POS=genomic position; REF=wild type nucleotide; ALT=mutated nucleotide; FREQ=percentage frequency in Morocco.

***Table S33: Mutations observed only in Sierra Leone and not in other parts of Africa***

| **POSITION** | **REF** | **ALT** | **AA CHANGE** | **REGION** | **NUMBER** | **FREQUENCY %** |
|---|---|---|---|---|---|---|

| | | | | | | |
|---|---|---|---|---|---|---|
| 10818 | C | T | A255V | *3C-like proteinase* | 3 | 27.3% |

POS=genomic position; REF=wild type nucleotide; ALT=mutated nucleotide; FREQ=percentage frequency in Africa.

### Proteins Affected

***Table S34: SARS CoV2 variants greater than 1% with associated frequencies, amino acid changes, and mutation type***

| POS | REF | ALT | % TOTAL | % MUT | %EGYPT | GENE/PROTEIN | AA CHANGE | TYPE | T/T |
|---|---|---|---|---|---|---|---|---|---|
| **3037** | C | T | 47.27% | 70.27% | 98.68% | *ORF1ab/nsp3* | F106F | Synonymous | Transition |
| **23403** | A | G | 47.27% | 70.27% | 98.24% | *S/surface glycoprotein* | D614G | Missense | Transition |
| **14408** | C | T | 47.27% | 70.27% | 30.40% | *ORF1ab/RdRp* | P323L | Missense | Transition |
| **241** | C | T | 45.45% | 67.57% | 95.15% | *5'-UTR* | NA | Non-coding | Transition |
| **25563** | G | T | 27.27% | 40.54% | 81.50% | *Orf3a* | Q57H | Missense | Transversion |
| **1059** | C | T | 18.18% | 27.03% | 0.44% | *ORF1ab/Nsp2* | T85I | Missense | Transition |
| **8782** | C | T | 12.73% | 18.92% | 0.44% | *ORF1ab/NSP4* | S2839S | Synonymous | Transition |
| **28144** | T | C | 12.73% | 18.92% | 0.00% | *ORF8* | L84S | Missense | Transition |
| **28878** | G | A | 12.73% | 18.92% | 0.00% | *N/nucleocapsid phosphoprotein* | S202K | Missense | Transition |
| **22468** | G | T | 10.91% | 16.22% | 0.44% | *S/ surface glycoprotein* | T302T | Synonymous | Transversion |
| **29742** | G | A | 10.91% | 16.22% | 0.00% | *3'-UTR Stem loop* | NA | Non-coding | Transition |
| **28881** | G | A | 9.09% | 13.51% | 8.37% | *N/nucleocapsid phosphoprotein* | R203K | Missense | Transition |
| **28882** | G | A | 9.09% | 13.51% | 8.37% | *N/nucleocapsid phosphoprotein* | R203K | Missense | Transition |
| **28883** | G | C | 9.09% | 13.51% | 8.37% | *N/nucleocapsid phosphoprotein* | G204R | Missense | Transversion |
| **18877** | C | T | 7.27% | 10.81% | 11.01% | *ORF1ab/3'-to-5' exonuclease* | L280L | Synonymous | Transition |
| **41** | T | C | 7.27% | 10.81% | 0.00% | *5'-UTR* | NA | Non-coding | Transition |
| **11083** | G | T | 7.27% | 10.81% | 0.00% | *ORF1ab/Nsp6* | L37F | Missense | Transversion |
| **15324** | C | T | 7.27% | 10.81% | 0.00% | *ORF1ab/RdRp* | N628N | Missense | Transition |
| **1684** | C | T | 5.45 | 8.11 | 0.44% | *ORF1ab/ nsp2* | I293I | Synonymous | Transition |
| **361** | A | G | 5.45 | 8.11 | 0.00% | *ORF1ab/leader protein* | G32G | Synonymous | Transition |
| **6404** | G | T | 5.45 | 8.11 | 0.00% | *ORF1ab/nsp3* | V1229F | Missense | Transversion |
| **8208** | C | T | 5.45 | 8.11 | 0.00% | *ORF1ab/nsp3* | T1830I | Missense | Transition |

| | | | | | | | | | |
|---|---|---|---|---|---|---|---|---|---|
| **8658** | A | G | 5.45 | 8.11 | 0.00% | *ORF1ab/nsp4* | K35R | Missense | Transition |
| **9802** | G | A | 5.45 | 8.11 | 0.00% | *ORF1ab/nsp4* | A416A | Synonymous | Transition |
| **10818** | C | T | 5.45 | 8.11 | 0.00% | *ORF1ab/3C-like proteinase* | A255V | Missense | Transition |
| **12970** | C | T | 5.45 | 8.11 | 0.00% | *ORF1ab/nsp10* | N95N | Synonymous | Transition |
| **16512** | A | G | 5.45 | 8.11 | 0.00% | *ORF1ab/helicase* | L92L | Synonymous | Transition |
| **19951** | C | T | 5.45 | 8.11 | 0.00% | *ORF1ab/ endoRNAse* | P111S | Missense | Transition |
| **22363** | G | T | 5.45 | 8.11 | 0.00% | *S/ surface glycoprotein* | V801V | Synonymous | Transversion |
| **26428** | G | T | 5.45 | 8.11 | 0.00% | *E/envelope protein* | V62F | Missense | Transition |
| **29362** | C | T | 5.45 | 8.11 | 0.00% | *N/ nucleocapsid phosphoprotein* | P364H | Missense | Transition |
| **40** | C | A | 3.64 | 5.41 | 0.00% | *5'-UTR* | NA | Non-coding | Transversion |
| **198** | G | A | 3.64 | 5.41 | 0.00% | *5'-UTR* | NA | Non-coding | Transition |
| **9214** | C | T | 3.64 | 5.41 | 0.00% | *ORF1ab/nsp4* | Y220Y | Synonymous | Transition |
| **10889** | C | T | 3.64 | 5.41 | 0.00% | *ORF1ab/3C-like proteinase* | R279C | Missense | Transition |
| **23127** | C | T | 3.64 | 5.41 | 0.00% | *S/ surface glycoprotein* | A522V | Missense | Transition |
| **25821** | C | T | 3.64 | 5.41 | 0.00% | *ORF3a/ ORF3a protein* | A143A | Synonymous | Transition |
| **29811** | T | C | 3.64 | 5.41 | 0.00% | *3'-UTR* | NA | Non-coding | Transition |
| **29813** | A | G | 3.64 | 5.41 | 0.00% | *3'-UTR* | NA | Non-coding | Transition |
| **29817** | T | G | 3.64 | 5.41 | 0.00% | *3'-UTR* | NA | Non-coding | Transversion |
| **29822** | T | C | 3.64 | 5.41 | 0.00% | *3'-UTR* | NA | Non-coding | Transition |
| **29824** | A | C | 3.64 | 5.41 | 0.00% | *3'-UTR* | NA | Non-coding | Transversion |
| **23593** | G | T | 1.82 | 2.70 | 3.96% | *S /Surface glycoprotein* | Q677H | Missense | Transversion |
| **2113** | C | T | 1.82 | 2.70 | 0.88% | *ORF1ab/nsp2* | I436I | Synonymous | Transition |
| **1404** | C | T | 1.82 | 2.70 | 0.44% | *ORF1ab/nsp2* | P200L | Missense | Transition |
| **10046** | G | A | 1.82 | 2.70 | 0.44% | *Nsp4* | V498I | Missense | Transition |
| **21575** | C | T | 1.82 | 2.70 | 0.44% | *S/ surface glycoprotein* | L5F | Missense | Transition |
| **6** | A | T | 1.82 | 2.70 | 0.00% | *5'-UTR* | NA | Non-coding | Transversion |
| **17** | T | C | 1.82 | 2.70 | 0.00% | *5'-UTR* | NA | Non-coding | Transition |
| **203** | C | T | 1.82 | 2.70 | 0.00% | *5'-UTR* | NA | Non-coding | Transition |
| **228** | C | T | 1.82 | 2.70 | 0.00% | *5'-UTR* | NA | Non-coding | Transition |

| | | | | | | | | | |
|---|---|---|---|---|---|---|---|---|---|
| **315** | G | T | 1.82 | 2.70 | 0.00% | *ORF1ab/leader protein* | S17I | Missense | Transversion |
| **391** | A | G | 1.82 | 2.70 | 0.00% | *ORF1ab/leader protein* | A42A | Synonymous | Transition |
| **745** | C | T | 1.82 | 2.70 | 0.00% | *ORF1ab/leader protein* | N160N | Synonymous | Transition |
| **1191** | C | T | 1.82 | 2.70 | 0.00% | *ORF1ab/nsp2* | P129L | Missense | Transition |
| **1397** | G | A | 1.82 | 2.70 | 0.00% | *ORF1ab/nsp2* | V198I | Missense | Transition |
| **1666** | T | C | 1.82 | 2.70 | 0.00% | *ORF1ab/nsp2* | F288F | Synonymous | Transition |
| **2306** | C | T | 1.82 | 2.70 | 0.00% | *ORF1ab/nsp2* | L501F | Missense | Transition |
| **2416** | C | T | 1.82 | 2.70 | 0.00% | *ORF1ab/nsp2* | Y537Y | Synonymous | Transition |
| **2480** | A | G | 1.82 | 2.70 | 0.00% | *ORF1ab/nsp2* | I559V | Missense | Transition |
| **2558** | C | T | 1.82 | 2.70 | 0.00% | *ORF1ab/nsp2* | P585S | Missense | Transition |
| **2113** | C | T | | | | *ORF1ab/nsp2* | I436I | Synonymous | Transition |
| **1404** | C | T | | | | *ORF1ab/nsp2* | P200L | Missense | Transition |
| **1059** | C | T | | | | *ORF1ab/nsp2* | T85I | Missense | Transition |
| **3340** | G | T | 1.82 | 2.70 | 0.00% | *ORF1ab/nsp3* | V207V | Synonymous | Transversion |
| **3896** | G | T | 1.82 | 2.70 | 0.00% | *ORF1ab/nsp3* | V393F | Missense | Transversion |
| **4288** | G | T | 1.82 | 2.70 | 0.00% | *ORF1ab/nsp3* | E523G | Missense | Transversion |
| **5359** | T | C | 1.82 | 2.70 | 0.00% | *ORF1ab/nsp3* | A880A | Synonymous | Transition |
| **5401** | A | G | 1.82 | 2.70 | 0.00% | *ORF1ab/nsp3* | A894A | Synonymous | Transition |
| **5557** | T | C | 1.82 | 2.70 | 0.00% | *ORF1ab/nsp3* | G946V | Missense | Transition |
| **5672** | C | T | 1.82 | 2.70 | 0.00% | *ORF1ab/nsp3* | P985S | Missense | Transition |
| **5986** | C | T | 1.82 | 2.70 | 0.00% | *ORF1ab/nsp3* | F1089F | Synonymous | Transition |
| **6040** | C | T | 1.82 | 2.70 | 0.00% | *ORF1ab/nsp3* | F1107F | Synonymous | Transition |
| **6312** | C | A | 1.82 | 2.70 | 0.00% | *ORF1ab/nsp3* | T1198L | Missense | Transversion |
| **6337** | G | T | 1.82 | 2.70 | 0.00% | *ORF1ab/nsp3* | S1206S | Synonymous | Transversion |
| **7279** | C | T | 1.82 | 2.70 | 0.00% | *ORF1ab/nsp3* | F1520F | Synonymous | Transition |
| **7420** | C | T | 1.82 | 2.70 | 0.00% | *ORF1ab/nsp3* | I1567I | Synonymous | Transition |
| **7765** | C | T | 1.82 | 2.70 | 0.00% | *ORF1ab/nsp3* | TCC->TCT S1682S | Synonymous | Transition |
| **8097** | C | T | 1.82 | 2.70 | 0.00% | *ORF1ab/nsp3* | ACA->ATA T1793I | Missense | Transition |
| **8371** | G | T | 1.82 | 2.70 | 0.00% | *ORF1ab/nsp3* | CAG->CAT Q1884H | Missense | Transversion |
| **8389** | C | T | 1.82 | 2.70 | 0.00% | *ORF1ab/nsp3* | AAC->AAT N1890N | Synonymous | Transition |

| | | | | | | | | | |
|---|---|---|---|---|---|---|---|---|---|
| **3373** | C | A | | | | *ORF1ab/nsp3* | D218E | Missense | Transversion |
| **4002** | C | T | | | | *ORF1ab/nsp3* | T428I | Missense | Transition |
| **5020** | C | T | | | | *ORF1ab/nsp3* | D767D | Synonymous | Transition |
| **3011** | T | C | | | | *ORF1ab/nsp3* | L98L | Synonymous | Transition |
| **4354** | G | T | | | | *ORF1ab/nsp3* | E545G | Missense | Transversion |
| **2841** | C | T | | | | *ORF1ab/nsp3* | A41V | Missense | Transition |
| **3037** | C | T | | | | *ORF1ab/nsp3* | F106F | Synonymous | Transition |
| **8655** | C | T | 1.82 | 2.70 | 0.00% | *ORF1ab/nsp4* | S34F | Missense | Transition |
| **8781** | G | A | 1.82 | 2.70 | 0.00% | *ORF1ab/nsp4* | S76N | Missense | Transition |
| **8950** | C | T | 1.82 | 2.70 | 0.00% | *ORF1ab/nsp4* | I132I | Synonymous | Transition |
| **9479** | G | T | 1.82 | 2.70 | 0.00% | *ORF1ab/nsp4* | G309C | Missense | Transversion |
| **9514** | A | G | 1.82 | 2.70 | 0.00% | *ORF1ab/nsp4* | L320L | Synonymous | Transition |
| **9483** | A | C | | | | *ORF1ab/nsp4* | Y220Y | Missense | Transversion |
| **9520** | C | T | | | | *ORF1ab/nsp4* | E310D | Missense | Transition |
| **10046** | G | A | | | | *ORF1ab/nsp4* | F322F | Synonymous | Transversion |
| **8782** | C | T | | | | *ORF1ab/nsp4* | V498I | Missense | Transition |
| **9483** | A | C | | | | *ORF1ab/nsp4* | S76S | Synonymous | Transversion |
| **10456** | C | T | 1.82 | 2.70 | 0.00% | *ORF1ab/3C-like proteinase* | F134F | Synonymous | Transition |
| **10582** | C | T | 1.82 | 2.70 | 0.00% | *ORF1ab/3C-like proteinase* | D176D | Synonymous | Transition |
| **10818** | C | T | | | | *ORF1ab/3C-like proteinase* | A255V | Missense | Transition |
| **10889** | C | T | | | | *ORF1ab/3C-like proteinase* | R279C | Missense | Transition |
| **10097** | G | A | | | | *ORF1ab/3C-like proteinase* | G15S | Synonymous | Transition |
| **11575** | C | T | 1.82 | 2.70 | 0.00% | *ORF1ab/nsp6* | F201F | Synonymous | Transition |
| **11782** | A | G | 1.82 | 2.70 | 0.00% | *ORF1ab /nsp6* | K270K | Synonymous | Transition |
| **11935** | A | C | 1.82 | 2.70 | 0.00% | *ORF1ab/nsp7* | Q31H | Missense | Transition |
| **13495** | C | T | 1.82 | 2.70 | 0.00% | *ORF1ab/stem-loop 1* | S21F | Missense | Transition |
| **13515** | C | T | 1.82 | 2.70 | 0.00% | *ORF1ab/RdRp* | G25G | Synonymous | Transition |
| **13517** | C | T | 1.82 | 2.70 | 0.00% | *ORF1ab/RdRp* | T26I | Missense | Transition |
| **13620** | C | T | 1.82 | 2.70 | 0.00% | *ORF1ab/RdRp* | D60D | Synonymous | Transition |
| **13730** | C | T | 1.82 | 2.70 | 0.00% | *ORF1ab/RdRp* | A97V | Missense | Transition |

| | | | | | | | | | |
|---|---|---|---|---|---|---|---|---|---|
| **13858** | G | T | 1.82 | 2.70 | 0.00% | *ORF1ab/RdRp* | D140Y | Missense | Transversion |
| **14184** | C | T | 1.82 | 2.70 | 0.00% | *ORF1ab/RdRp* | T248T | Synonymous | Transition |
| **14805** | C | T | 1.82 | 2.70 | 0.00% | *ORF1ab/RdRp* | Y455Y | Synonymous | Transition |
| **15372** | G | T | 1.82 | 2.70 | 0.00% | *ORF1ab/RdRp* | T644T | Synonymous | Transversion |
| **15380** | G | T | 1.82 | 2.70 | 0.00% | *ORF1ab/RdRp* | S647I | Missense | Transversion |
| **16178** | C | T | 1.82 | 2.70 | 0.00% | *ORF1ab/RdRp* | S913L | Missense | Transition |
| **13536** | C | T | | | | *ORF1ab/RdRp* | Y32Y | Synonymous | Transition |
| **14408** | C | T | | | | *ORF1ab/RdRp* | P323L | Missense | Transition |
| **15324** | C | T | | | | *ORF1ab/RdRp* | N628N | Synonymous | Transition |
| **14805** | C | T | | | | *ORF1ab/RdRp* | Y455Y | Synonymous | Transition |
| **16575** | C | T | 1.82 | 2.70 | 0.00% | *Helicase* | | | Transition |
| **17690** | C | T | 1.82 | 2.70 | 0.00% | *Helicase* | | | Transition |
| **18508** | C | T | 1.82 | 2.70 | 0.00% | *3'-to-5' exonuclease* | L157F | Missense | Transition |
| **18928** | C | T | 1.82 | 2.70 | 0.00% | *3'-to-5' exonuclease* | P297S | Missense | Transition |
| **19525** | G | T | 1.82 | 2.70 | 0.00% | *3'-to-5' exonuclease* | D496Y | Missense | Transversion |
| **19735** | G | T | 1.82 | 2.70 | 0.00% | *endoRNAse* | | | Transversion |
| **20032** | C | T | 1.82 | 2.70 | 0.00% | *endoRNAse* | | | Transition |
| **20374** | A | G | 1.82 | 2.70 | 0.00% | *endoRNAse* | | | Transition |
| **20759** | C | T | 1.82 | 2.70 | 0.00% | *2'-O-ribose methyltransferase* | | | Transition |
| **21595** | C | T | 1.82 | 2.70 | 0.00% | *S/ surface glycoprotein* | V11V | Missense | Transition |
| **21648** | C | T | 1.82 | 2.70 | 0.00% | *S/ surface glycoprotein* | T29I | Missense | Transition |
| **22210** | C | T | 1.82 | 2.70 | 0.00% | *S/ surface glycoprotein* | L216L | Synonymous | Transition |
| **22397** | T | A | 1.82 | 2.70 | 0.00% | *S/ surface glycoprotein* | Y279N | Missense | Transversion |
| **22424** | C | T | 1.82 | 2.70 | 0.00% | *S/ surface glycoprotein* | A288V | Missense | Transition |
| **22435** | T | C | 1.82 | 2.70 | 0.00% | *S/ surface glycoprotein* | C291R | Missense | Transition |
| **22503** | A | G | 1.82 | 2.70 | 0.00% | *S/ surface glycoprotein* | Q314Q | Synonymous | Transition |
| **23923** | C | T | 1.82 | 2.70 | 0.00% | *S/ surface glycoprotein* | Q787H | Missense | Transition |
| **24370** | C | T | 1.82 | 2.70 | 0.00% | *S/ surface glycoprotein* | S937L | Missense | Transition |
| **24926** | G | T | 1.82 | 2.70 | 0.00% | *S/ surface glycoprotein* | V1122V | Synonymous | Transversion |
| **24992** | G | C | 1.82 | 2.70 | 0.00% | *S/ surface glycoprotein* | E1144E | Synonymous | Transition |

| | | | | | | | | | |
|---|---|---|---|---|---|---|---|---|---|
| **25217** | G | T | 1.82 | 2.70 | 0.00% | *S/ surface glycoprotein* | G1219C | Missense | Transversion |
| **23403** | A | G | | | | *S/ surface glycoprotein* | D614G | Missense | Transition |
| **23127** | C | T | | | | *S/ surface glycoprotein* | A522V | Missense | Transition |
| **21575** | C | T | | | | *S/ surface glycoprotein* | TTT L5F | Missense | Transition |
| **22468** | G | T | | | | *S/ surface glycoprotein* | T302T | Synonymous | Transversion |
| **25411** | A | C | 1.82 | 2.70 | 0.00% | *ORF3a/ ORF3a protein* | I7L | Missense | Transversion |
| **25702** | C | T | 1.82 | 2.70 | 0.00% | *ORF3a/ ORF3a protein* | P104S | Missense | Transition |
| **26144** | G | T | 1.82 | 2.70 | 0.00% | *ORF3a/ ORF3a protein* | G251V | Synonymous | Transversion |
| **25563** | G | T | | | | *ORF3a/ ORF3a protein* | Q57H | Missense | Transversion |
| **25821** | C | T | | | | *ORF3a/ ORF3a protein* | A143A | Missense | Transition |
| **26299** | C | T | 1.82 | 2.70 | 0.00% | *E/Envelope protein* | L19F | Missense | Transition |
| **26428** | G | T | | | | *E/Envelope protein* | V62F | Missense | Transversion |
| **26257** | G | T | | | | *E/Envelope protein* | V5F | Missense | Transversion |
| **26665** | T | C | 1.82 | 2.70 | 0.00% | *M/membrane glycoprotein* | I48T | Missense | Transition |
| **26720** | G | C | 1.82 | 2.70 | 0.00% | *M/membrane glycoprotein* | V66V | Synonymous | Transversion |
| **26775** | G | T | 1.82 | 2.70 | 0.00% | *M/membrane glycoprotein* | A85S | Missense | Transversion |
| **26790** | T | C | 1.82 | 2.70 | 0.00% | *M/membrane glycoprotein* | L90L | Synonymous | Transition |
| **26801** | C | G | 1.82 | 2.70 | 0.00% | *M/membrane glycoprotein* | L93L | Synonymous | Transversion |
| **27520** | A | T | 1.82 | 2.70 | 0.00% | *ORF7a/ORF7a protein* | N43Y | Missense | Transition |
| **27754** | G | T | 1.82 | 2.70 | 0.00% | *ORF7a/ORF7aprotein* | Z121END | **Nonsense** | Transversion |
| **27769** | C | T | 1.82 | 2.70 | 0.00% | *ORF7b/ QOI10401.1* | S5L | Missense | Transition |
| **27804** | C | T | 1.82 | 2.70 | 0.00% | *ORF7b/ QOI10401.1* | L17L | Synonymous | Transition |
| **28878** | G | A | | | | *N/nucleocapsid phosphoprotein* | S202N | | |
| **28881** | G | A | | | | *N/nucleocapsid phosphoprotein* | R203K | | |
| **28882** | G | A | | | | *N/nucleocapsid phosphoprotein* | R203R | | |
| **28883** | G | C | | | | *N/nucleocapsid phosphoprotein* | G204R | | |

| | | | | | | | | | |
|---|---|---|---|---|---|---|---|---|---|
| **28311** | C | T | 1.82 | 2.70 | 0.00% | *N/nucleocapsid phosphoprotein* | P13L | Missense | Transition |
| **28346** | G | T | 1.82 | 2.70 | 0.00% | *N/ nucleocapsid phosphoprotein* | G25C | Missense | Transversion |
| **28347** | G | T | 1.82 | 2.70 | 0.00% | *N/ nucleocapsid phosphoprotein* | G25V | Missense | Transversion |
| **28531** | C | T | 1.82 | 2.70 | 0.00% | *N/ nucleocapsid phosphoprotein* | Y86Y | Synonymous | Transition |
| **28688** | T | C | 1.82 | 2.70 | 0.00% | *N/ nucleocapsid phosphoprotein* | L139L | Synonymous | Transition |
| **28738** | T | A | 1.82 | 2.70 | 0.00% | *N/ nucleocapsid phosphoprotein* | A155A | Synonymous | Transition |
| **28830** | C | A | 1.82 | 2.70 | 0.00% | *N/ nucleocapsid phosphoprotein* | S186Y | Missense | Transversion |
| **28835** | T | C | 1.82 | 2.70 | 0.00% | *N/ nucleocapsid phosphoprotein* | S188P | Missense | Transition |
| **28854** | C | T | 1.82 | 2.70 | 0.00% | *N/ nucleocapsid phosphoprotein* | S194L | Missense | Transition |
| **28985** | G | T | 1.82 | 2.70 | 0.00% | *N/ nucleocapsid phosphoprotein* | G238C | Missense | Transversion |
| **29315** | G | C | 1.82 | 2.70 | 0.00% | *N/ nucleocapsid phosphoprotein* | D348H | Missense | Transversion |
| **29449** | G | T | 1.82 | 2.70 | 0.00% | *N/ nucleocapsid phosphoprotein* | V393V | Missense | Transversion |
| **29474** | G | T | 1.82 | 2.70 | 0.00% | *N/ nucleocapsid phosphoprotein* | D401Y | Missense | Transversion |
| **29748** | T | C | 1.82 | 2.70 | 0.00% | *S2m* | Noncoding | NA | Transition |
| **14362** | C | T | 0.00% | 0.00% | 50.71% | *ORF1ab/RdRp* | F307F | Synonymous | Transition |
| **29871** | A | G | 0.00% | 0.00% | 50.71% | *3' UTR* | Noncoding | NA | Transition |
| **15907** | G | A | 0.00% | 0.00% | 50.35% | *ORF1ab/RdRp* | Q822K | Synonymous | Transition |
| **17091** | T | C | 0.00% | 0.00% | 50.35% | *ORF1ab/helicase* | G285G | Synonymous | Transition |
| **26257** | G | T | 0.00% | 0.00% | 50.35% | *E/Envelope protein* | V5F | Missense- | Transversion |
| **29744** | G | A | 0.00% | 0.00% | 4.26% | *3'UTR* | NA | Non-coding | Transition |

| | | | | | | | | | |
|---|---|---|---|---|---|---|---|---|---|
| **28849** | C | T | 0.00% | 0.00% | 3.90% | *N/ nucleocapsid phosphoprotein* | N192N | Synonymous | Transition |
| **5284** | C | T | 0.00% | 0.00% | 3.19% | *ORF1ab/nsp3* | N855N | Synonymous | Transition |
| **10097** | G | A | 0.00% | 0.00% | 3.19% | *ORF1ab/ 3C-like proteinase* | G15S | Missense | Transition |
| **12534** | C | T | 0.00% | 0.00% | 3.19% | *ORF1ab/nsp8* | T148I | Synonymous | Transition |
| **28908** | G | T | 0.00% | 0.00% | 3.19% | *N/ nucleocapsid phosphoprotein* | G212V | Missense | Transversion |
| **23731** | C | T | 0.00% | 0.00% | 2.84% | *S/surface glycoprotein* | T723T | Synonymous | Transition |
| **3373** | C | A | 0.00% | 0.00% | 2.48% | *ORF1ab/nsp3* | D218E | Missense | Transversion |
| **4002** | C | T | 0.00% | 0.00% | 2.48% | *ORF1ab/nsp3* | ACT->ATT T428I | Missense | Transition |
| **13536** | C | T | 0.00% | 0.00% | 2.48% | *ORF1ab/RdRp* | ACA->ATA T4424I | Missense | Transition |
| **5020** | C | T | 0.00% | 0.00% | 2.13% | *ORF1ab/nsp3* | D767D | Synonymous | Transition |
| **3011** | T | C | 0.00% | 0.00% | 1.77% | *ORF1ab/nsp3* | L98L | Synonymous | Transition |
| **4354** | G | T | 0.00% | 0.00% | 1.77% | *ORF1ab/nsp3* | E545G | Missense | Transversion |
| **28846** | C | T | 0.00% | 0.00% | 1.77% | *N/ nucleocapsid phosphoprotein* | R191R | Synonymous | Transition |
| **2841** | C | T | 0.00% | 0.00% | 1.06% | *ORF1ab/nsp3* | A41V | Missense | Transition |
| **9483** | A | C | 0.00% | 0.00% | 1.06% | *ORF1ab/nsp4* | E310D | Missense | Transversion |
| **9520** | C | T | 0.00% | 0.00% | 1.06% | *ORF1ab/nsp4* | F322F | Synonymous | Transition |
| **11991** | A | G | 0.00% | 0.00% | 1.06% | *ORF1ab/nsp7* | E50G | Missense | Transition |
| **17550** | C | T | 0.00% | 0.00% | 1.06% | *ORF1ab/helicase* | L438L | Missense | Transition |
| **17766** | C | T | 0.00% | 0.00% | 1.06% | *ORF1ab/helicase* | V510V | Missense | Transition |
| **21597** | C | T | 0.00% | 0.00% | 1.06% | *S/surface glycoprotein* | S12F | Missense | Transition |

POS=genomic position; REF=wild type nucleotide; ALT=mutated nucleotide; %TOTAL=percent frequency in all samples excluding Egypt; AA CHANGE = amino acid change; T/T= transition or transversion.
